# Approximate Bayesian computation untangles signatures of contemporary and historical hybridization between two endangered species

**DOI:** 10.1101/2021.02.24.432626

**Authors:** Hannes Dittberner, Aurelien Tellier, Juliette de Meaux

## Abstract

Contemporary gene flow, when resumed after a period of isolation, can have crucial consequences for endangered species, as it can both increase the supply of adaptive alleles and erode local adaptation. Determining the history of gene flow and thus the importance of contemporary hybridization, however, is notoriously difficult. Here, we focus on two endangered plant species, *Arabis nemorensis* and *A. sagittata*, which hybridize naturally in a sympatric population located on the banks of the Rhine. Using reduced genome sequencing, we determined the phylogeography of the two taxa but report only a unique sympatric population. Molecular variation in chloroplast DNA indicated that *A. sagittata* is the principal receiver of gene flow. Applying classical D-statistics and its derivatives to whole-genome data of 35 accessions, we detect gene flow not only in the sympatric population but also among allopatric populations. Using an Approximate Bayesian computation approach, we identify the model that best describes the history of gene flow between these taxa. This model shows that low levels of gene flow have persisted long after speciation. Around 10 000 years ago, gene flow stopped and a period of complete isolation began. Eventually, a hotspot of contemporary hybridization was formed in the unique sympatric population. Occasional sympatry may have helped protect these lineages from extinction in spite of their extremely low diversity.

## INTRODUCTION

Individual taxa do not always evolve in isolation. Interspecific hybridization, when it leads to fertile offspring, can result in the transfer of alleles across species barriers and even allow to speed the pace of adaptation (Seehausen 2004; Servedio et al. 2013, Abbott 2017, Nieto Feliner et al. 2017, Todesco et al. 2020). The footprints of gene flow are detectable in many genera and demes (Seehausen 2004; Marcet-Houben and Gabaldón 2015; Ackermann et al. 2019; Taylor and Larson 2019). Genomic analyses have revealed that species barriers are established progressively, depending on the size and degree of reproductive isolation of the species, with genetic variation being shared over periods of time that are longer than previously thought (Brandvain et al. 2014; Novikova et al. 2016; Edelman et al. 2019; Small et al. 2020). The climatic oscillations of quaternary glaciation cycles have likely contributed to multiple opportunities for hybridization in many taxa (Hewitt 2000).

The evolutionary consequences of hybridization can be manifold, offering a powerful channel for some taxa to capture and quickly fix alleles that have been subjected to selection in other species (Hedrick 2013; Goulet et al. 2017; Suarez-Gonzalez et al. 2018a; Vallejo Marín and Hiscock 2018). The potent adaptive potential unleashed by hybridization have been confirmed in a number of species (Rieseberg et al. 2003; Baduel et al. 2018; Ma et al. 2019; Marburger et al. 2019). For example, introgression appeared to accelerate local adaptation via the transfer of alleles contributing to the success of a relative of the sunflower (Todesco et al. 2020). In *Heliconius* butterflies, an inversion associating genes that control color patterns has been repeatedly exchanged between species (Edelman et al. 2019). In addition, transgressive phenotypic variation can emerge from the recombination of resident and incoming alleles throughout the genome (Rieseberg et al. 1999; Seehausen 2004). Positive selection is therefore expected to increase introgression rates specifically in and around adaptive loci. Heterogeneous introgression rates, however, can also arise along the genome in the absence of adaptive gene flow (Schumer, Rosenthal, et al. 2018). Covariation between the rate of introgression, recombination and gene density, for example, indicates that selections acts to limit introgression and points to the existence of polygenic barriers to gene flow (Brandvain et al. 2014; Schumer, Xu, et al. 2018). In humans, the impact of deleterious alleles was shown to correlate negatively with the frequency of alleles introgressed from the related species *Homo neanderthalis,* whose population size was low and eventually collapsed (Juric et al. 2016; Steinrücken et al. 2018). Ultimately, high rates of gene flow can introduce maladapted alleles and ultimately impede adaptation (Lenormand 2002; Yeaman 2015; Tigano and Friesen 2016). These alleles, in turn, can select for alleles that will reinforce species barriers to prevent resources from being wasted by producing poorly performing hybrids (Hopkins and Rausher 2012). Gene flow, regardless of whether it promotes or erodes adaptation, is expected to create a heterogeneous pattern of introgression throughout the genome (Martin and Jiggins 2017; Schumer, Rosenthal, et al. 2018).

If gene flow can be either a blessing or a curse, it is particularly crucial to understand its importance for species that have to be protected from extinction. In a context of global climate change, the number of threatened species is expected to increase (Díaz et al. 2019; Eichenberg et al. 2020). At the same time, many species barriers that have so far been maintained by habitat, phenological or behavioral separation are likely to disintegrate, generating unprecedented opportunities for novel episodes of hybridization (Anderson et al. 2012; Chunco 2014). Contemporary hybridization will most affect endangered species if the taxa involved have been separated long enough to acquire distinct ecological and population genomics characteristics. For example, the hybridization of long separated continental sub-species of salmon has been associated with the manifestation of Dobzhansky-Muller incompatibilities (Rougemont and Bernatchez 2018). In *Mimulus*, barriers to gene flow are strong today despite an ancient history of hybridization, but contemporary hybridization has been reported at specific locations (Brandvain et al. 2014; Kenney and Sweigart 2016). The potential of hybridization to create a novel genetic make-up today that may impact future evolution and support the evolutionary rescue of biodiversity depends, therefore, on how much time has elapsed during the period of isolation.

Determining the history of gene flow between taxa is a complex task. Dating gene flow cannot be achieved with a single summary statistic of genomic variation, because effective population sizes, migration rates and/or the time since species formation jointly influence patterns of standing variation, divergence and expected allele sharing. The widely used D-statistics tests for gene flow, while accounting for incomplete lineage sorting at the time of speciation (Durand et al. 2011). This and related statistics have, for example, revealed the extent of gene flow between species thought to be separated by differences in ploidy (Arnold et al. 2016; Paape et al. 2018; Kryvokhyzha et al. 2019). Yet such statistics do not allow speciation or gene flow to be dated, nor do they evaluate the duration of isolation periods (Schumer, Rosenthal, et al. 2018; Hibbins and Hahn 2019). Failing to understand the correct history of speciation and gene flow can lead to confounding neutral patterns of gene flow with convergent evolution, sympatric speciation or even adaptive divergence (Bierne et al. 2013; Ravinet et al. 2017).

The evolutionary and demographic histories of hybridizing species are often too complex to be determined analytically. Additional population parameters such as population size and recombination rates have also been shown to have strong consequences on the amount of native genomic DNA that can be rescued (Harris et al. 2019). Intensive simulation methods, such as Approximate Bayesian Computation (ABC), offer a powerful alternative (Csilléry et al. 2010). It not only makes it possible to choose among competing models for the one best able to explain the data, but also enables population parameter (population sizes, migration rates) and their fluctuation over time to be estimated (Roux et al. 2013; Leroy et al. 2017; Fraïsse et al. 2018; Rougemont and Bernatchez 2018). However, ABC approaches have rarely been applied to whole-genome data and when, then only to infer relatively simple demographic events such as bottlenecks or expansions (Boitard et al. 2016, Jay et al. 2019). The current limitation stems from the inherent statistical complexity of the approach (high dimensionality of whole-genome data), the necessity for an educated guess regarding the choice of summary statistics to use, and the lack of user-friendly ready-to-use software capable of handling all genome data. The potential of ABC approaches to determine the timing, intensity and duration of hybridization episodes based on whole genome data thus remains to be fully leveraged.

Here, we focused on a documented case of contemporary hybridization between two endangered plant species and reconstructed the history of gene flow. The species we examined, *Arabis nemorensis* and *A. sagittata,* are selfing biennial forbs from the Brassicaceae family. *A. nemorensis*, which is strictly confined to floodplain environments in Central and Northern Europe, harbors extremely low levels of nucleotide diversity, with about 1.5 SNP expected in 10kb (Dittberner et al. 2019). Since the 1950s, conversion to arable land and management intensification have reduced the area covered by species–rich floodplain meadows, the ecosystem in which *A. nemorensis* thrives, by more than 80 %. Only small remnant patches have persisted within protected areas (Hölzel 2005). The unique ecology of *A. nemorensis*, its shrinking habitat and its selfing mating system all contribute to the acute danger of extinction this species finds itself in (Hölzel 2005; Burmeier et al. 2010; Mathar et al. 2015). *A. nemorensis*, however, was found to occur in sympatry with its relative, *A. sagittata*, in a small set of pristine habitat patches located on the banks of the Rhine near Mainz, Germany (Dittberner et al. 2019). *A. sagittata*, known to thrive in relatively dry environments, is also endangered, but its presence in floodplain meadows is novel (Hand and Gregor 2006; Dittberner et al. 2019). Interestingly, approximately 10% of the individuals of the sympatric population appeared to be of mixed ancestry (Dittberner et al. 2019).

Using a combination of reduced sequencing and whole-genome sequencing, we asked the following questions: Is hybridization local or is there a large contact zone? Is gene flow symmetrical? Do rates of introgression vary along the genome? Is hybridization contemporary or was gene flow continuous throughout the history of the species? Our study confirms the existence of a contemporary hotspot of hybridization where asymmetric gene flow has resumed after approximately 10 000 years of complete isolation between taxa.

## RESULTS

### Admixed individuals detected only in a single sympatric population

We combined previously published and newly generated RAD-sequencing data for a total of 231 accessions to describe the phylogeographic distribution of *A. nemorensis* and assess the distribution of *A. sagittata* and admixed individuals in the area of distribution of *A. nemorensis* (Table S1). This collection of genotypes covered all sites where the presence of *A. nemorensis* had been documented (see methods). We performed an admixture analysis (Alexander and Lange 2011a) and identified two genetic clusters, that were previously assigned to *A. nemorensis* and *A. sagittata* (Figure S1-5, Dittberner et al. 2019). We found individuals of the two species growing in sympatry in only one of the eleven sites (the Rhine population), indicating that opportunities for natural hybridization are restricted (Figure 1A, Dittberner et al. 2019). Of the 140 individuals sampled in the sympatric population, 75 were *A. sagittata*, 42 were *A. nemorensis* and 23 were individuals we identified as having mixed ancestry because they showed less than 95% purity in the output of the ADMIXTURE analysis. Phenotypic observations in the field, such as low seed number per silique and elongated stems, had suggested that hybridization may occur in some rare instances (Novotná and Czapik 1974; Titz 1979). In contrast, genetic analyses indicate that the two species frequently cross-hybridize within the sympatric population (Dittberner et al. 2019). Interestingly, *A. sagittata* ancestry predominates in most admixed individuals (Figure 1B), indicating the frequent backcross of admixed with *A. sagittata* individuals, which are more frequent than *A.nemorensis* individuals in the sympatric population.

**Figure 1:**
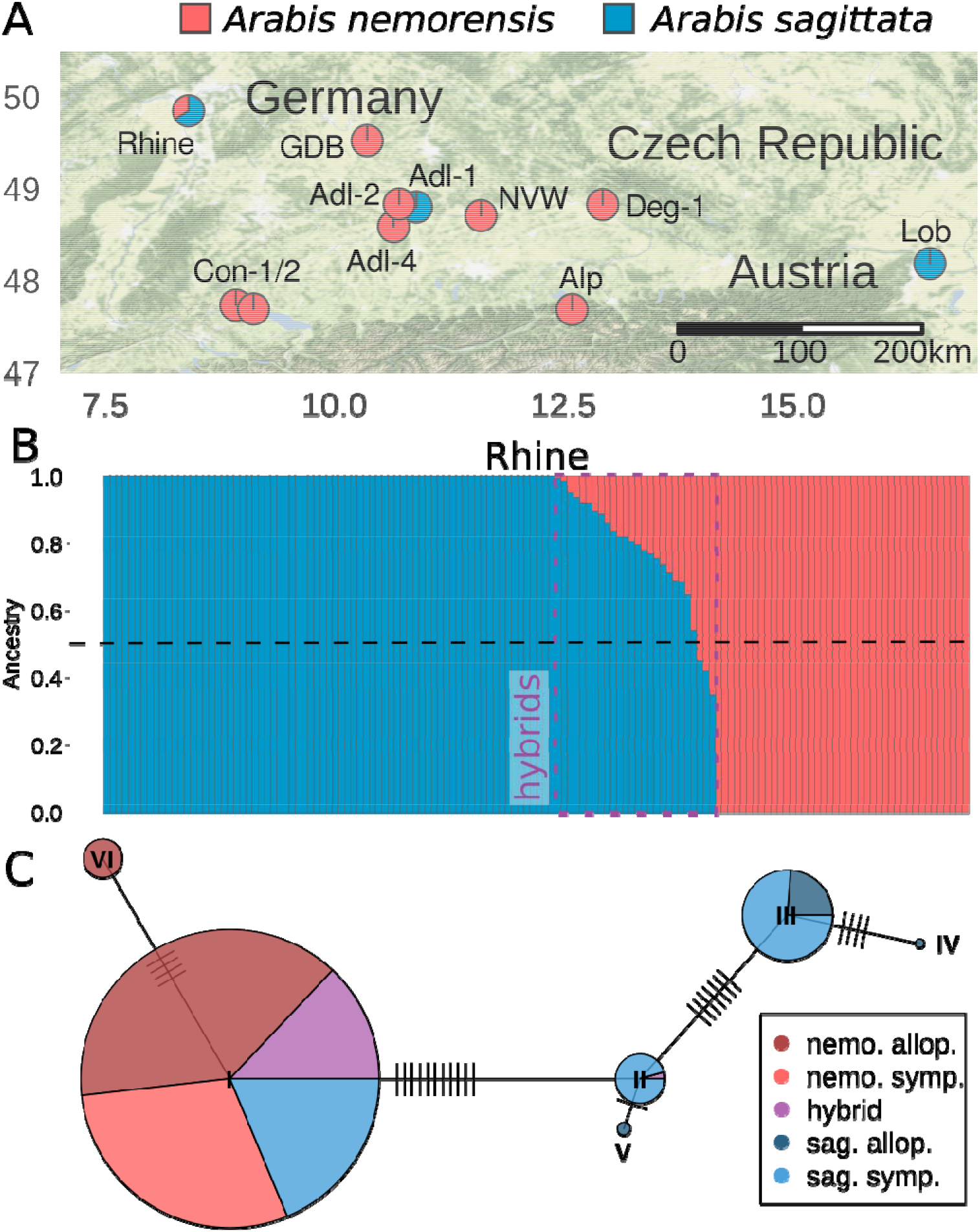
Hybridization between A. nemorensis and A. sagittata. A) Map of sampled populations. Each population is represented by a pie chart showing the average ancestry proportions of A. nemorensis and A. sagittata in the given population, based on RAD-sequencing data. B) Representation of individual ancestry components in the sympatric (“Rhine”) population, based on RAD-seq data. Each bar represents one individual and is colored according to its genomic ancestry. Bars are ordered by decreasing A. sagittata ancestry. Admixed individuals are framed with a purple rectangle. C) Network of chloroplast RAD-seq haplotypes. Each pie-chart represents one haplotype and the fractions of species/populations carrying this haplotype. Haplotypes are connected to their closest relative by a line. Orthogonal dashes represent the number of mutations. Abbreviations in the legend: *nemo. - A. nemorensis, sag. - A. sagittata, allo. - allopatric, symp. - sympatric*.

### Chloroplast DNA indicates that *A. nemorensis* is the maternal parent of hybrids

Chloroplast DNA is maternally inherited and its sequence variation can provide information about the maternal genotype of hybrids (McCauley 1995). To determine the maternal genotypes of hybrids, we focused on RADseq stacks that mapped to a total of 10 kb (5%) of the chloroplast sequence of the close relative *Arabis hirsuta* (Kawabe et al. 2018). Haplotype network analysis showed that the chloroplast sequences of *A. sagittata* formed four closely related haplotype groups and two for *A. nemorensis* (Figure 1C). As all but one admixed individual had an *A. nemorensis* haplotype, we conclude that *A. nemorensis* is the maternal genotype of most hybrids. This observation confirms that gene flow between the two clusters is asymmetrical. Surprisingly, 24 *A. sagittata* individuals were found to carry the *A. nemorensis* chloroplast haplotype although the admixture analysis found no trace of *A. nemorensis* in their nuclear genome. This suggests that some individuals may have a hybrid ancestry but their *A. nemorensis* ancestry is undetectable in the admixture analysis of RAD-seq data, showing in turn that the examination of whole-genome sequences was necessary.

### Gene flow between *A. nemorensis* and *A. sagittata* is not restricted to the sympatric population

To quantify interspecific gene flow between the two genetic clusters, we resequenced 35 whole genomes of accessions from both allopatric and sympatric populations for the two species, as well as one individual of the closest diploid relative, *A. androsacea* (Table S2). In order to understand the history of gene flow, we specifically excluded individuals that were predicted to be admixed (Fig. 1B), which were obviously formed a handful of generations ago. Analyses presented hereafter were all conducted on data generated by whole-genome sequencing in this set of 35 accessions. We confirm that nucleotide diversity is low in this system (at synonymous sites, π= 1.32 e-5 and 4.37e-5 in *A. sagittata* and *A. nemorensis*, respectively). The two species are clearly differentiated with a median Fst = 0.8, yet median Dxy is 0.0003 and net divergence Da = 0.03%, which lie in the range of values observed between populations of the same species that have the potential to exchange gene flow (Roux et al. 2016). We first computed Patterson’s *D* (ABBA-BABA) statistic (Durand et al. 2011; Green et al. 2010), a statistic that quantifies gene flow after accounting for incomplete allele sorting. This statistic is computed over the whole genome for phylogenies of the form (((P1,P2),P3),O), where O is the outgroup *A. androsacea* and P1 and P2 two populations of the same species. D was highest for the phylogeny with P1 = *A. sagittata* allopatric, P2 = *A. sagittata* sympatric and P3 = all *A. nemorensis* individuals with 0.2 (p<<0.001;block jackknife test). This significant and positive D value shows that gene flow is higher between P3 *A. nemorensis* and P2, the sympatric *A. sagittata* population, than between *A. nemorensis* and P1, the allopatric *A. sagittata* individuals (Figure 2A). Although this test showed elevated rates of gene flow in the sympatric population, it did not rule out introgressions in the allopatric populations. Thus, we also compared the two allopatric *A. sagittata* populations Lob and Adl-1 (Figure 1A) with P1 = *A. sagittata* Lob, P2 = *A. sagittata* Adl-1 and P3 = all *A. nemorensis*, and found *D* was 0.09 (p<<0.001; block jackknife test) indicating significant gene flow outside of the sympatric population. Finally, we tested the phylogeny with P1 = *A. nemorensis* sympatric, P2 = *A. nemorensis* allopatric and P3 = *A. sagittata* Lob (allopatric), and found that *D* was 0.17 (p<<0.001, block jackknife test). This positive value indicated that gene flow from *A. sagittata* into *A. nemorensis* was stronger in allopatry than in sympatry, demonstrating that the history of hybridization in this system predates the formation of the sympatric population.

**Figure 2:**
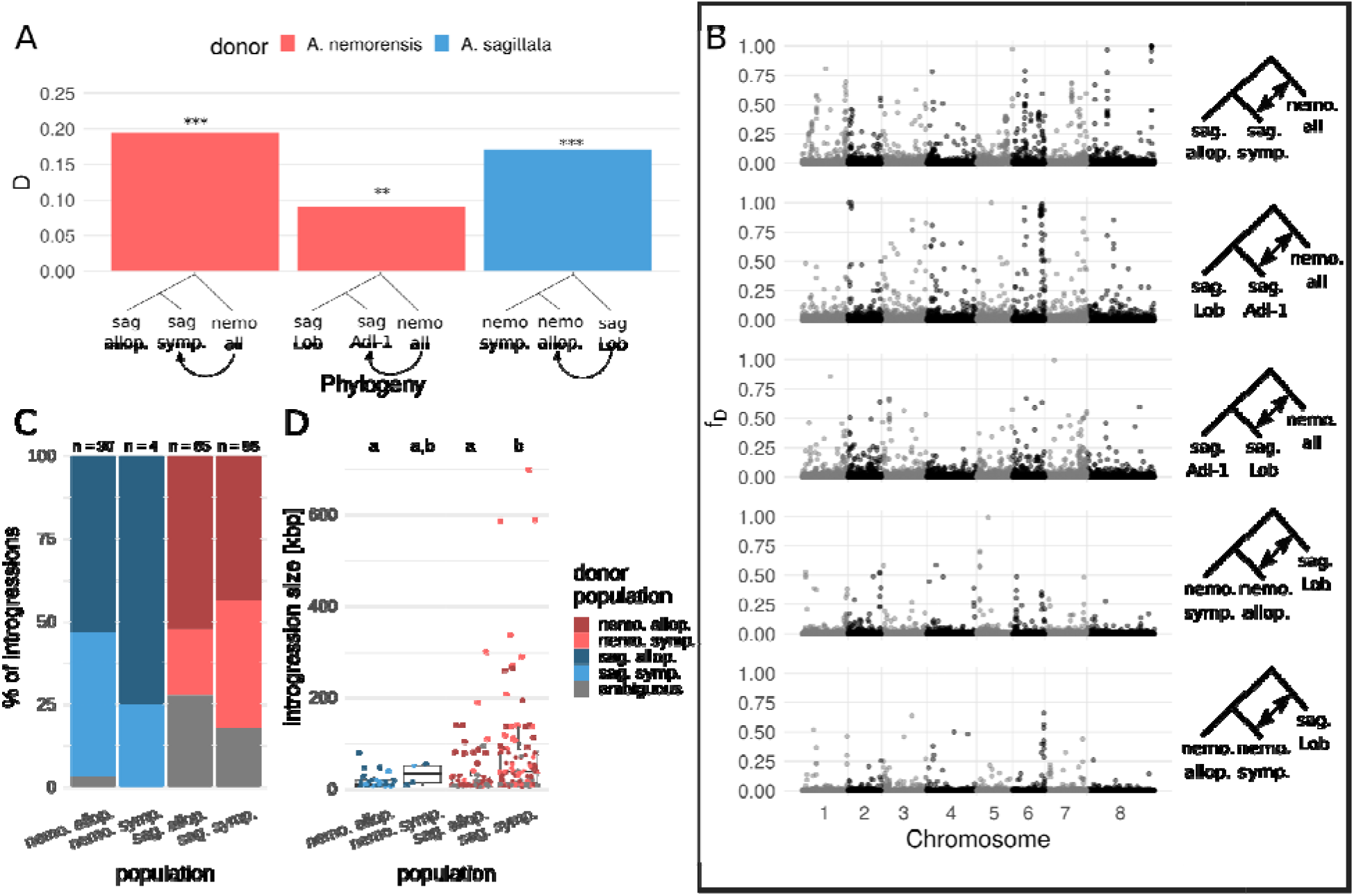
Patterns of introgression in sympatric and allopatric populations of parental species. A) D-statistic results calculated over the whole genome for different phylogenies. Bar color represents the species of the P3 / donor population. Asterisks represent the result of a Jackknife test: *** p<0.001, ** p < 0.1 B) Genome-wide distribution of fD calculated in 50-kbp windows for different phylogenies, as shown on the right. Plots are ordered by decreasing overall genome-wide fD. C) Fractions of introgression origins for each population, inferred based on minimum genetic distances (see Methods). Numbers on top show the total number of introgressions in each population. This plot does not take into account the frequency of the identified introgression tracts. D) Distribution of introgression size in each population, represented by dots and boxplots. Dots are colored according to the introgression origin. This plot does not take into account the frequency of the identified introgression tracts. The compact letter display indicates significant differences among groups. Abbreviations in the legends: *nemo. - A. nemorensis, sag. - A. sagittata, allo. - allopatric, symp. - sympatric*

### The frequency of introgression is heterogeneous along the genome

The *D*-statistic provides information about genome-wide rates of gene flow but not about its distribution across the genome. Thus, we calculated the f_D_-statistic (Martin et al. 2015) across the genome for five phylogenies covering all possible scenarios of interspecific gene flow (Figure 2B). Distributions of f_D_ were highly zero-inflated for all populations, indicating that introgressions were generally rare across the genome. Yet, in all populations, we found genomic regions with elevated f_D_ values. We did not find strong differences in the distribution of these regions among chromosomes. The D and f_D_ statistics rely on the assumption that there was no gene flow between P3 and P1 (Green et al. 2010; Durand et al. 2011; Martin et al. 2015). As we found introgressions in all populations, this assumption was violated, meaning any introgression shared by P1 and P2 would remain undetected. Our estimates of gene flow are therefore likely to be conservative.

### Introgressed fragments are on average largest in the sympatric population

To locate introgressed fragments and their boundaries in the genome, we calculated the ratio of average intra- and interspecific genetic distance for each individual in 10 kb genomic windows (see Methods). This ratio is expected to be smaller than the ratio throughout the genome, except in introgressed regions. We used a value equal or greater than 2 as a conservative threshold for the ratio value calling introgressed fragments. The average number of introgressions per individual was highest in the sympatric *A. sagittata* population, with 24.7 introgressed fragments per genome (standard deviation (sd) = 3.6), followed by the allopatric *A. sagittata* populations, with an overall mean of 13.7 fragments per genome (sd = 3.5, Table S3). In contrast, in *A. nemorensis,* we observed an average of 9.6 introgressed fragments per genome (sd = 1.47) in the sympatric population and 2.6 (sd = 0.55) in the allopatric. These results confirmed that interspecific gene flow from *A. nemorensis* to *A. sagittata* was stronger than vice versa. Furthermore, the presence of introgressions in allopatric populations suggests that gene flow is at least partly historical.

The high occurrence of introgression fragments in the sympatric *A. sagittata* population indicates that some of these fragments could have been introduced more recently by contemporary gene flow. We thus assessed whether the introgression was likely to have occurred in the sympatric population (recent gene flow) or whether it predated the separation of local populations (historical gene flow). For this, we determined the individual in the donor species, whose orthologous region was most closely related to the introgressed fragment identified in the receiver species. We then observed that the proportion of introgressed fragments that were most similar to alleles of the sympatric *A. nemorensis* population was almost twice as high in the sympatric *A. sagittata* population (38.5%) as in the allopatric *A. sagittata* population (20%), a difference that was marginally significant (Chi square= 3.76, p=0.05, Figure 2C). This trend was reversed in *A. nemorensis,* with 25% of introgressions originating from a sympatric *A. sagittata* lineage in the sympatric *A. nemorensis* population as opposed to 43% in the allopatric *A. sagittata* population. However, as there were only four introgressions in the sympatric population, this reversal is likely due to chance.

We further reasoned that recent introgressions originating from sympatric donor lineages should be on average larger than introgressions originating from allopatric donor lineages because the latter being more ancient, they should have had more time to be broken down by recombination. Thus, we compared the distributions of introgression size among populations. In *A. sagittata*, introgressions in the sympatric population were significantly larger than in the allopatric population (Z=-4.47, p<0.001), with median values of 37,900 bp and 10,000 bp, respectively (Figure 2D). We did not find this difference in the *A. sagittata* allopatric populations (Kruskal-Wallis chi-squared = 0.5174, df = 1, p = 0.472) or in the two *A. nemorensis* populations (symp.: Kruskal-Wallis chi-squared = 0.6, df = 1, p = 0.4386; allop.: Kruskal-Wallis chi-squared = 1.772, df = 1, p = 0.1831).

### The best fit model excludes constant gene flow between the two taxa

Based on the above, we observed i) asymmetrical gene flow between species, ii) evidence for gene flow both within and outside of the sympatric population. Our data thus indicate that gene flow may have been continuous throughout the history of the two species. However, we also observed that gene flow from *A. nemorensis* into *A. sagittata* might have intensified in the sympatric population. Gene flow could thus have occurred in the past and stopped during a phase of complete isolation, only to resume recently in the sympatric population. We took a modelling ABC approach to date gene flow between *A. nemorensis* and *A. sagittata* and to estimate its strength and past fluctuation.

We modelled the history of interspecific gene flow using a random-forest-based approximate Bayesian computation (ABC) approach (Raynal et al. 2019). Our goal was to determine whether the data is explained better by a history of continuous or by episodic gene flow. We thus chose five demographic models, which explored different modes of intra- and interspecific gene flow (Figure 3A), and generated 50,000 coalescent simulations under each one. In the first model we assumed no migration at all, while symmetric intraspecific migration was assumed in all other models. In the second model, interspecific migration stopped 100,000 generations after the species split. In the third model ancestral interspecific migration continued until the first intraspecific population split. The fourth model was an extension of the third model that additionally allowed migration between the sympatric populations, after populations in both species had split. In the fifth model, interspecific migration continued throughout the history of the species, allowing a change in intensity after the intraspecific population had split in both species. All interspecific migration rates were allowed to be asymmetrical. Population sizes were constant for each population but were allowed to change at all population split points.

**Figure 3:**
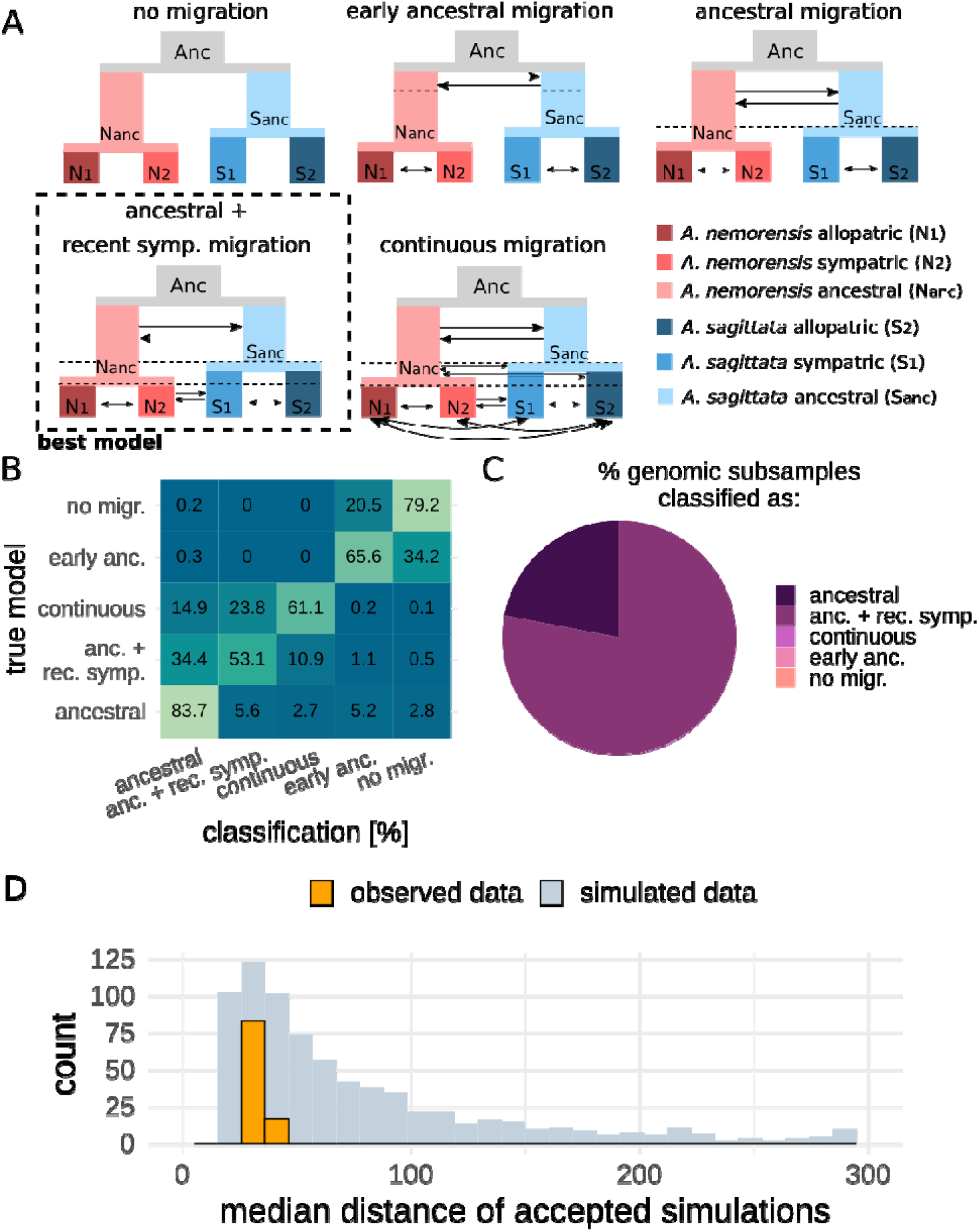
Choosing the best demographic model. A) Schematic representation of the tested demographic models. Populations are colored according to the bottom-right code. Arrows represent migration rates. Interspecific migration rates can be asymmetric (even with double-sided arrows) and intraspecific migration is symmetric. Dashed lines are timepoints at which populations split and/or migration rates change. B) Confusion matrix of the model-choice random forest model. For each model, simulated (out-of-bag) samples were classified by the trained random forest. Correct classifications are on the diagonal. Results are represented a percentages. C) Mean model classification results, i.e. random forest votes, for 100 observed datasets, with 500 randomly selected genomic windows each. D) Evaluation of fit for the best model. For each of the 100 observed datasets, the median normalized summary statistic distance between observed data and the closest 1% of simulated datasets was calculated (orange). As a null distribution, the same calculation was done with 1000 simulated datasets of pseudo-observed data. For better clarity, the x-axis was trimmed at 300, but no observed distance was larger than that.

To choose the model that best fitted the data, we first trained a random-forest classifier on the data all five models use. The accuracy of the trained classifier was evaluated with a confusion matrix that described how many simulations were assigned to the model under which they were generated (Figure 3B). Based on this matrix, we identified two subsets of models that summary statistics could accurately distinguish (max. error 5.2 %). The first subset comprised the two models with the least migration: the model without any migration and the model with early ancestral interspecific migration. The second subset contained three models with one or more episodes of interspecific migration: prolonged ancestral migration, ancestral and recent sympatric migration separated by a period of complete isolation, and continuous migration. Within this last subset, the first model was classified with a high accuracy of 84 %, but the second and third models were only classified with accuracies of 53 % and 61 %, respectively. These results show that i) models with both high and low migration rates could be distinguished easily by the classifier; and ii) determining the exact model within each of the two groups was less straightforward. This result was not surprising because when recent migration is low, the ancestral and recent migration model gives results similar to those generated under the ancestral migration model, which does not allow migration in the sympatric population.

Next, we used the trained random-forest classifier to determine the model that best explained the observed data. To this end, we randomly selected 100 datasets, each consisting of 500 loci of individual length 75 kbp sampled from the observed genotype (full genome) data and summarized with 200 summary statistics (Table S4). Models with non-constant gene flow always received the majority of the votes. The model with ancestral & recent sympatric migration was chosen as the best model for 78% of the observed genomic subsamples (Figure 3C). This was followed by the model assuming ancestral migration only for 22 % of observed genomic subsamples. The other three models were not selected as the most likely model.

Ancestral and recent sympatric migration separated by a period of isolation most aptly characterize the history of this plant system (Figure 3A). A history of low but uninterrupted gene flow (Model 5) can be clearly ruled out. We further quantified the goodness-of-fit of the model we identified as the best. The distance between observed and simulated datasets was not significantly different from zero, confirming that the observed data lay well within expectations of the model (Figure 3D). Next, we used this model to estimate the timing, strength and direction of gene flow between species and populations as well as its fluctuation across genome sub-samples.

### Estimation of demographic parameters indicates very low Ne in both taxa

To estimate demographic parameters of the best model, we generated a total of 360,000 coalescent simulations under this demographic model. We used two methods for parameter estimation -- random-forest-based ABC (Marin et al. 2019), which also generated confidence intervals, and extreme gradient boosting (XGBoost; Chen et al. 2020), which provided the most accurate point estimates -- and estimated each parameter independently (see Methods for details). We achieved highest accuracy for the estimation of present population sizes and lowest accuracy for the time and population size at speciation (Figure S6-7). Of the two methods, XGBoost tended to have the lowest RMSE and the highest R2, indicating that it could best reconstruct the parameters under which simulated data was generated (Figure S8).

We then estimated the demographic parameters of the two species based on observed data. We present in detail the estimates obtained for one of the observed datasets we randomly picked among those assigned to the selected model. The estimated contemporary population sizes for *A. nemorensis* were 6,775 (sd = 1,988) for the allopatric and 2,964 (sd = 615) for the sympatric population (Figure 4, Table S5). For *A. sagittata*, 14,895 (sd = 2,407) made up the effective allopatric population and 6,425 (sd = 1,649), the effective sympatric population. These results agree with those of the previous report, namely, that in the sympatric population, *A. sagittata* harbored levels of genetic diversity higher than those of *A. nemorensis* (Dittberner et al. 2019). The sizes estimated for the allopatric populations were larger than the estimated size of the sympatric populations in both species, presumably because the allopatric samples included individuals collected in several sites. We further estimated that the ancestral population in *A. sagittata* was more than twice as large as that in *A. nemorensis*, with 59,031 (sd = 12,529) and 25,206 (sd = 9,777), respectively. We note that this result also indicates that a simple scenario of early speciation followed by secondary contact is unlikely to explain the data. The magnitude of the population size decline after the split in both species, with a mean factor of 6.11 for *A. nemorensis*, which was similar to the factor of 6.57 estimated for *A. sagittata*. A decline of effective population sizes in the recent past has also been observed in the closely related species *Arabidopsis thaliana,* which is also selfing (Durvasula et al. 2017). However, contemporary population sizes in this species are still larger by at least an order of magnitude. This difference indicates that *A. nemorensis* and *A. sagittata* have not been particularly successful in their natural landscape.

**Figure 4:**
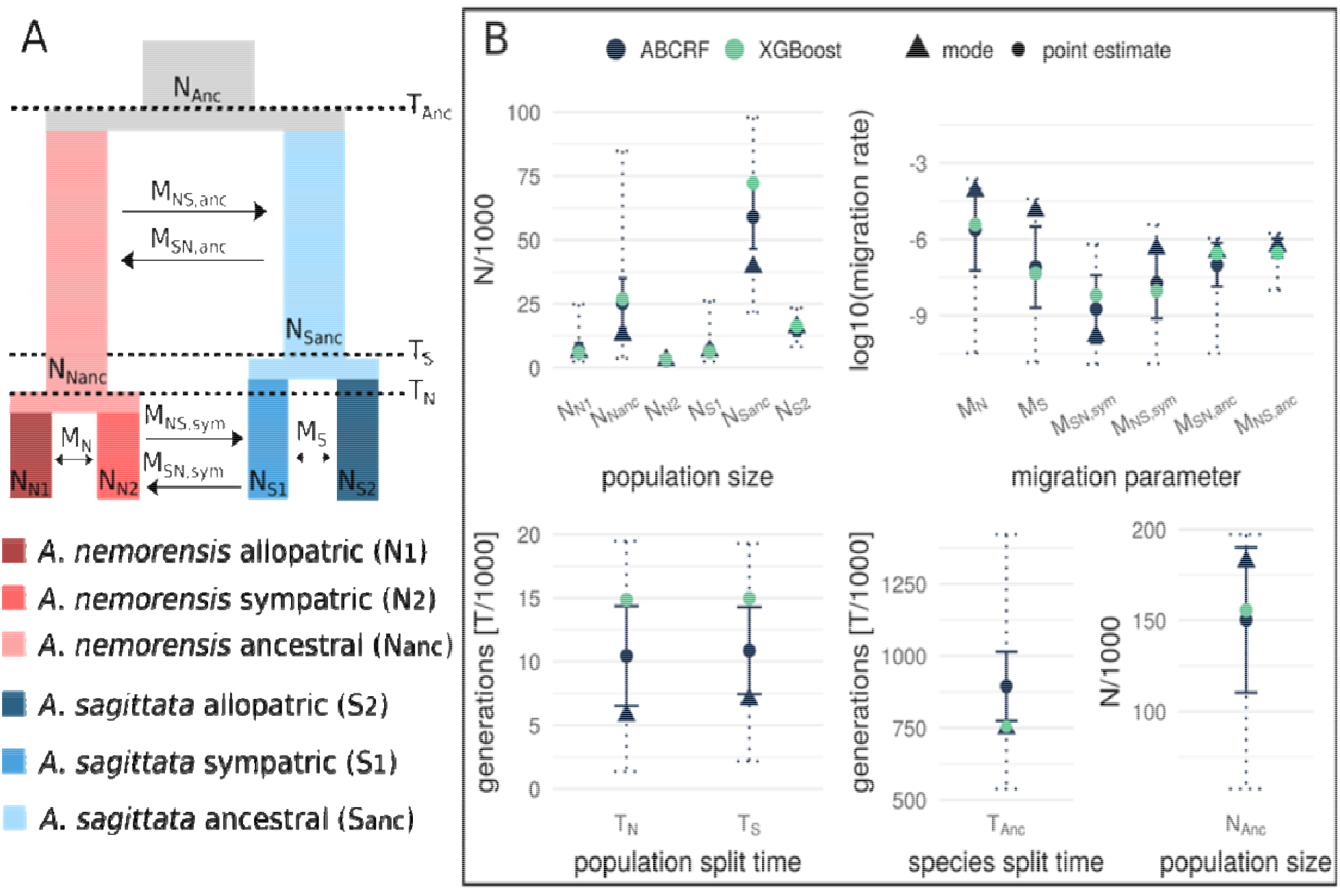
Model parameter estimation. A) Schematic representation of the model and its parameters: N = effective population size, M = migration rate (fraction of migrants per generation), T = Species/population split times. B) Estimation results of the model parameters based on two methods: ABCRF and XGBoost. Points represent the point estimate for each method, and triangles, the mode of the posterior distribution estimated by ABCRF. Solid error bars represent the standard deviation of the point estimate of ABCRF. Dashed error bars represent the 95% prediction interval of the ABCRF posterior distribution.

### The period of isolation dates back to the last glaciation

The split between the species occurred approximately 894,801 (sd = 119,548) generations ago, when the effective size of the ancestral population was approximately 150,281 (sd = 39,960). Standard deviations were high, presumably because our model also allowed for migration between populations (Figure S7). Within species, the populations split almost simultaneously: 10,441 (sd = 3,910) generations ago in *A. nemorensis* and 10,858 (sd = 3,417) generations ago in *A. sagittata*, and their split time coincides with the last glacial maximum. Our analysis thus suggests that the genetic landscape of the two taxa was established after the last glaciation, a period during which they became completely isolated.

### Estimates of migration rate confirmed gene flow resumed in sympatry

The low but significant estimates of gene flow showed that the model with ancestral and recent migration explained the data better than a model assuming only ancestral migration (Figure 3). We report the migration rates as the log10-transformed fraction of a population migrating per generation. Migration from *A. nemorensis* to *A. sagittata* in the ancestral population was approximately five times higher than vice versa: −6.34 (sd = 0.36) and −7.00 (sd = 0.85), respectively. This amounts to about one migrant every 33 generations. Point estimates indicated that, in the sympatric population, migration rates were lower than in the ancestral population. Yet, the rate was still approximately ten times higher from *A. nemorensis* to *A. sagittata* than vice versa, with a rate of −7.72 (sd = 1.39) and −8.74 (sd = 1.33). The mode of the posterior distribution for these two parameters differed even more strongly (−6.36 for migration from *A. nemorensis* to *A. sagittata* and −9.80 for the opposite direction). This result aligns well with the asymmetry of gene flow revealed by the analysis of chloroplast genome variation. As expected estimates of intraspecific migration rate were higher than interspecific migration rates. We noted that the estimate was two orders of magnitude higher in *A. nemorensis* compared to *A. sagittata* (−5.61, sd = 1.61 and −7.09, sd = 1.60 in *A. nemorensis* and *A. sagittata*, respectively).

### Variation in the observed genomic sample affects ancestral population size and sympatric migration rates

Since we observed that the introgression rate varies across the genome, we examined whether estimates of gene flow varied depending on the genomic regions included in the observed sample. To test this, we compared the distributions of random-forest-based and xgboost-based point estimates of all demographic parameters for the 100 observed datasets (Table S6). Estimates were fairly robust to genome subsamples because they were always in the same order of magnitude (Figure 5).

**Figure 5:**
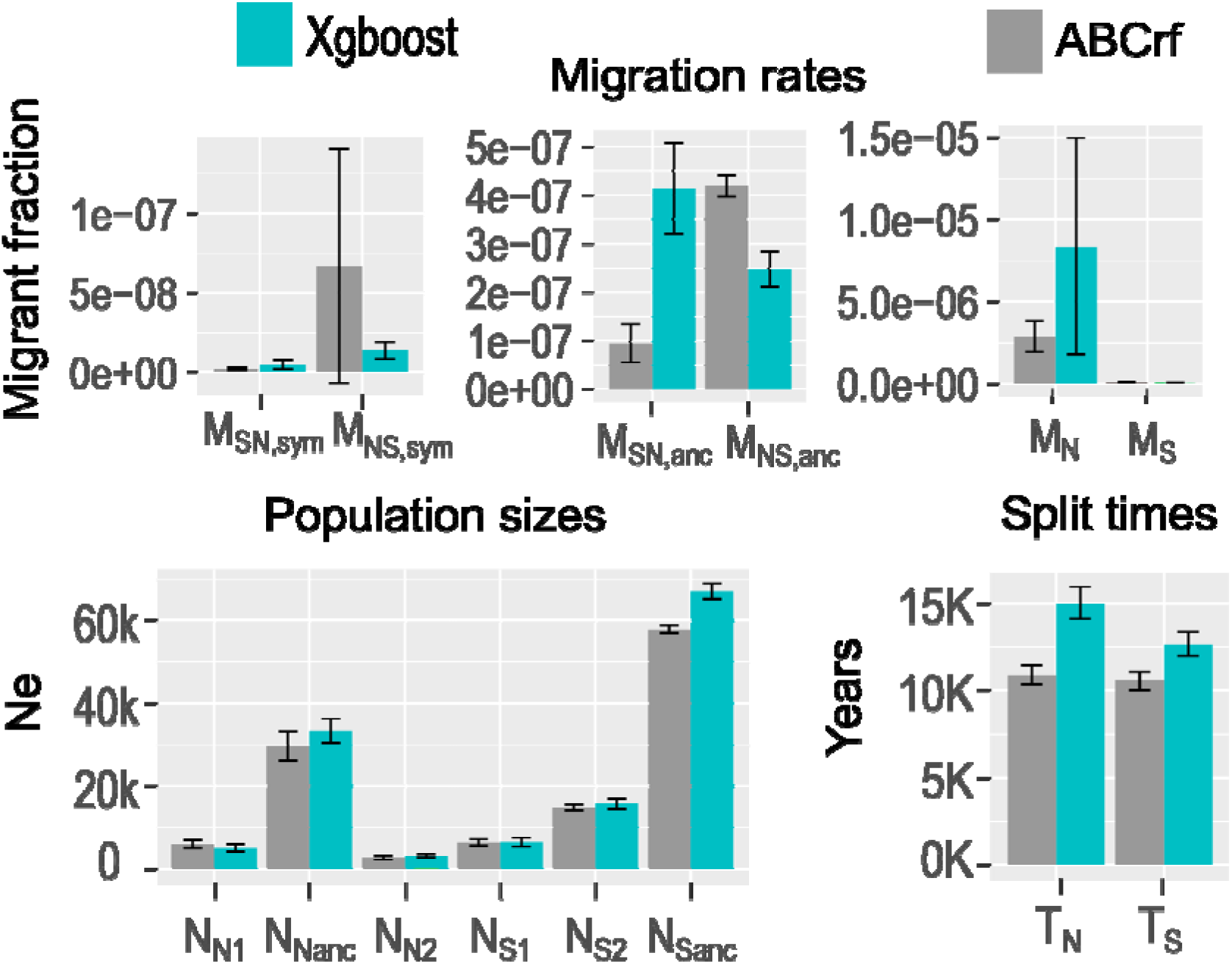
Distribution of parameter estimates for the 100 observed datasets formed by sub-sampling 500 75kb-long regions in the genome. M_N_ and M_S_: intraspecific migration rate among A. nemorensis and A. sagittata populations, respectively. M_NS_ and M_SN_: interspecific migration rate from A. nemorensis into A. sagittata, respectively. Sym: sympatric population, Anc. : ancient gene flow. N_N1_ : Effective population size of allopatric A. nemorensis populations, N_N2_ : Effective population of sympatric A. nemorensis population, N_S1_: Effective population of sympatric A. sagittata population, N_S2_– Effective population size of allopatric A. sagittata population, N_Nanc_: Ancestral population size of A. nemorensis N_Nanc_, N_Sanc_ : Ancestral population size of A. sagittata, T_N_ : time elapsed since split of A. nemorensis population, T_S_ : Time elapsed since split of A. sagittata populations, T_Anc_: time of speciation, N_Anc_ : ancestral population size prior to speciation.

Yet, we made two notable observations. First, the abcrf estimate of sympatric migration from *A. nemorensis* to *A. sagittata* was 4.6 times more variable than the estimate of migration in the opposite direction (Figure 5). The distribution of point estimates for recent sympatric migration from *A. nemorensis* to *A. sagittata* ranged from −8.3 to −6.5, with a mean of −7.5 (Figure S9). Since the estimate was positively correlated with the proportion of introgression regions included in each genomic sub-sample, it partly reflects the heterogeneity of introgression rates along the genome (Spearman correlation coefficient Rho=0.2, p= 0.03). The estimate of local introgression rate fd also showed a weak yet significantly negative relationship with inferred recombination rates in the genome (Spearman Rho = −0.06, p= 0.005, Figure S10) indicating that introgressed fragments were more likely in less recombining regions of the genome. However, we note that gradient boosting, a machine learning method that does not take the variance of the coalescent process into account, yielded migration estimates that were little affected by the composition of the genomic loci sampled for each analysis. Second, while both methods indicated the same asymmetry in gene flow between *A. nemorensis* and *A. sagittata* in the sympatric population, they yielded opposite conclusions for ancestral gene flow. Drawing a firm conclusion on the direction of ancestral gene flow is therefore not possible.

## DISCUSSION

### ABC modelling untangles contemporary and ancestral gene flow between two taxa

Explorations of both genome-wide and local estimates of interspecific introgressions, as well as allelic distributions within and between species at introgressed loci, indicated that these two Arabis species share a history of gene flow that predates the formation of the sympatric population. The potential of ABC approaches had been revealed previously by the study of population isolation in *Salmo salar* (Rougemont and Bernatchez 2018). Patterns of gene flow in *S. salar* populations, however, were resolved by genotyping more than 2,000 individuals at some 5,000 SNPs (Rougemont and Bernatchez 2018). This study now shows that ABC approaches have the power to disentangle complex demographic scenarios even in endangered species, where large sample of genotypes are by definition not available. Indeed, we provide compelling evidence that, although low levels of gene flow were maintained long after speciation, interspecific gene flow was not continuous in the history of these two plant species. Models postulating a phase of complete isolation initiated approximately 10,000 generations ago, were markedly better at explaining the data than those assuming constant gene flow.

Several aspects could potentially challenge this conclusion. First, gene flow may be underestimated, if many incompatibilities or deleterious alleles had to be removed by natural selection (Roux et al. 2016, Sousa et al. 2013). Second, our model ignored the effect of linked selection, which, by reducing Ne locally, can bias demographic inferences and decrease effective local recombination (Shrider et al. 2016). In both cases, a higher rate of introgression would be expected in regions that are recombining more actively (Schumer, Xu et al. 2018). The data, however, indicated the opposite, with a marginally higher introgression rates in low recombining regions of the genome. Third, the allopatric populations were grouped into a single metapopulation, whereby some aspect of demographic complexity could have been ignored. However, none of these simplifications could lead to caveats that would explain why, in the sympatric population of *A. sagittata*, we observed longer introgressed fragments than in the allopatric population. The migration rates estimated for past gene flow were low in absolute number but they fit well the model in which gene flow has resumed only a some rare sympatric populations. We note that the estimate of gene flow in the sympatric population should be considered as the lower bound since the sample used for ABC demographic inference excluded individuals with obvious signatures of admixture. Intuitively, we believe that gene flow might have happened in the past as it happens today: through the occasional formation of sympatric demes leading to localized bursts of interspecific gene flow.

### Interspecific gene flow as a chance for evolutionary rescue?

*A. nemorensis* and *A. sagittata* are both considered endangered in Germany and in many parts of Europe (Dittberner et al. 2019). The demographic model confirms that effective population sizes are particularly small in these two low-diversity taxa (Figure S2-S5). Genetic diversity in both species was approximately an order of magnitude lower than in more common selfing relatives, such as e.g. *Arabidopsis thaliana* (Alonso-Blanco et al. 2016) or *Capsella rubella* (Onge et al. 2011). These low levels of diversity, however, do not necessarily point to a low adaptive potential that would increase their risk of extinction. Indeed, the hotspot of contemporary gene flow we uncovered has formed after a period of isolation that might have been sufficiently long to allow functional divergence. Variance in abcrf estimates associated with the sub-sampling of genomic variation indicates that the signature of contemporary gene flow is not homogeneous throughout the genome, in contrast to that of ancient gene flow. Such a heterogeneity suggests that the contemporary contact is not without fitness consequences, and, indeed, we found several high-frequency introgressions that could be a signature of adaptive gene flow (Racimo et al. 2017). Studies in organisms such as yeast or sunflower have shown that hybrid populations can adapt efficiently to changing environmental conditions (Todesco et al. 2020; Mitchell et al. 2019; Stelkens et al. 2014). Moreover, gene flow can also resolve fitness tradeoffs limiting adaptation in the parental species (Walter et al. 2020).

At this stage, however, we don’t know whether, in this system, heterogeneous gene flow along the genome reflects positive selection for introgression (Suarez-Gonzalez et al. 2018; Shumer et al. 2018). First, we do not know whether the two species possess alleles of potential fitness relevance for each other. Second, F1 hybrids display low fertility, indicating that gene flow causes fertility reduction, at least initially (HD pers. com.; Titz 1979). The simple removal of a handful of large effect incompatibility alleles could contribute to the pattern of heterogeneous introgression documented here (Schumer et al. 2014). Introgressions may also carry deleterious alleles, either due to fitness trade-offs between the parental species (Arnegard et al. 2014) or due to slightly deleterious variants in the parental genomes that have not been purged by selection, i.e., genetic load. This phenomenon is known to have shaped the landscape of introgression from Neanderthals to humans (Harris and Nielsen 2016; Juric et al. 2016), resulting in more introgressions outside of either functionally important regions or of regions with low recombination rates (Schumer, Xu, et al. 2018). The absence of a global positive correlation between recombination rate and introgression suggests that negative selection is likely weak and not pervasive throughout the genome. Alternatively, transient heterozygosity following hybridization may favor gene flow in low recombining regions because these regions presumably harbor more deleterious variants that could be masked by the introgression. Novel methods for the detection of adaptive gene flow have been proposed, but this task remains arduous in selfing species, where recombination events are rare (Setter et al. 2020). Experimental work is required to identify the phenotypic consequences of gene flow between *A. sagittata* and *A. nemorensis* and disentangle the effect of positive and negative selection on the pattern of introgression. Having identified a local hotspot of contemporary hybridization, it is now possible to monitor these consequences in situ.

## MATERIAL AND METHODS

### Plant material and DNA extraction

In 2016 and 2017, we identified 30 sites reported to host *A. nemorensis* populations on the Deutschland Flora Database. Of these, 24 could be visited, and *A. nemorensis/A. sagittata* populations were observed in 11 of them. We sampled seeds from at least 10 *A. nemorensis* and/or *A. sagittata* plants per site for a total of 231 accessions (Table S1). Populations were located in Southern Germany and Austria (Figure 1A). One of these sites, referred to as “Rhine,” was previously described in (Dittberner et al. 2019) and consists of multiple proximate pristine habitat patches at one site. Individuals from these patches were assembled in one sympatric population. We extracted DNA for genotyping as previously described (Dittberner et al. 2019).

### RAD-seq genotyping for phylogeography

In addition to the 140 accessions originating from the Rhine populations that were previously genotyped in Dittberner et al. (2019), we genotyped 91 accessions using the original RAD-seq protocol (Etter et al. 2011) with the modification described in (Dittberner et al. 2019). Libraries were sequenced at the Cologne Center for Genomics on three Illumina HiSeq 4000 lanes with 2 × 150bp. We used *FastQC* (Andrews 2010) to check the raw reads. We trimmed adapters and removed reads shorter than 100bp using *Cutadapt* (Martin 2011). We removed PCR duplicates based on a 5 bp stretch of random nucleotides at the end of the adapter, using the *clone_filter* module of *Stacks* version 1.37 (Catchen et al. 2013). We de-multiplexed samples using the *process_radtags* module from *Stacks*. We filtered reads with ambiguous barcodes (allowed distance 2) and cut-sites, reads with uncalled bases and low-quality reads (default threshold).

We used our previously described pipeline to call nuclear genotypes in all samples (Dittberner et al. 2019). Briefly, we used BWA (Li and Durbin 2009) to map reads against the high-quality reference genome we had sequenced and assembled for *A. nemorensis* (Dittberner et al. 2019). We filtered mapped reads using *SAMtools* (Li et al. 2009) to remove regions with excessively low (30%) or high (2-fold) coverage, compared to the mean coverage. We called genotypes using samtools mpileup and VarScan2 (Koboldt et al. 2012). We filtered genotyped loci using VCFtools (Danecek et al. 2011). Knowing that both species are predominantly selfing, we filtered out SNPs with more than 20% overall heterozygosity and SNPs with more than 75% heterozygosity within a population, because heterozygous SNPs tended to cluster on a few single RAD-seq fragments, indicating inaccuracies in mapping that were not picked up by filtering on coverage. In total, we genotyped 2.6 million sites (∼ 1% of the genome) (excluding sites with more than 5% missing data in our sample) and identified 25,634 single nucleotide polymorphisms (SNPs).

To obtain SNPs in the chloroplast sequence, we mapped the RAD-seq reads against a chloroplast reference genome of *Arabis hirsuta* (Kawabe et al. 2018), using the *mem* algorithm of BWA (Li and Durbin 2009) with default settings. We called genotypes and SNPs as described above. We set heterozygous genotype calls to missing data, as these calls most likely resulted from mapping errors (the plastome is effectively haploid). Furthermore, we used VCFtools to remove SNPs with either more than 20% missing data or more than two alleles, which resulted in 23 chloroplast SNPs.

### Determining admixed individuals with RAD-seq and chloroplast data

To determine the genomic make-up of each population and identify the presence of admixed genotypes, we first analyzed SNP data using ADMIXTURE (Durand et al. 2011), varying K from 2 to 10. First we converted VCF files to bed format using PLINK (Purcell et al. 2007; Purcell 2009). We ran ADMIXTURE analysis (Alexander and Lange 2011a) for K=1 to K=6, with 10 iterations of cross-validation each. We normalized clusters across runs using CLUMPAK (Kopelman et al. 2015). We used K=2 for further analysis, as we were analyzing two species and the value was well supported by the cross-validation error (Figure S1). Individuals were defined as admixed if they had less than 95% ancestry from either species. We created plots using the libraries *ggplot2* (Wickham 2009), *ggmap* (Kahle and Wickham 2013), *scatterpie* (Yu 2018) and *ggsn* (Baquero 2017). Finally, we used the library *pegas* (Paradis et al. 2016b), to determine chloroplast haplotypes and build a haplotype network.

### Analysis of genetic diversity

We used the RAD-seq data to determine the level of genetic diversity within and between species. Using the vcfR package (Knaus and Grünwald 2017), the genotype data were loaded into R and converted to DNAbin format. We used the *pegas* package (Paradis et al. 2016b) to calculate pairwise genetic distances among all individuals. Based on the resulting distance matrix, we calculated average genetic distances within and between both populations and species. We calculated the diversity in the hybrid complex as the average genetic distance within the whole Rhine population.

### Whole genome resequencing for gene flow quantification and ABC

To achieve greater resolution for introgression detection, we randomly selected five *A. nemorensis* and 14 *A. sagittata* accessions from the sympatric population, 4 and 6 accessions from two allopatric *A. sagittata* populations and one accession of each of the six allopatric *A. nemorensis* populations. In total, the whole genome of 35 accessions was sequenced (Table S2). To provide an outgroup species for estimating gene flow, we also sequenced one accession of *Arabis androsacea*, provided by Jean-Gabriel Valay (Jardin Alpin du Lautaret, France). DNA for these accessions was extracted as described above. Libraries were prepared using Illumina TruSeq DNA PCR-free kits at the Cologne Center for Genomics. Six samples were sequenced on a HiSeq 4000 with 50 million 2×75 bp reads per sample. The remaining samples were sequenced on a NovaSeq6000 with 25 million 2×150 bp reads per sample, resulting in a depth of ∼ 25x.

We used *FastQC* (Andrews 2010) to quality-check the resulting reads. We filtered the reads using the *process_shortreads* module from *Stacks* v2.2 (Catchen et al. 2013), removing reads shorter than 70 bp or 100 bp (threshold was adapted to the sequencing device) and reads with uncalled bases or low-quality scores (default threshold). At this point, we included reads of the *A. nemorensis* reference genome accession from the sympatric population (Dittberner et al. 2019). We mapped the reads against the *A. nemorensis* reference genome (Dittberner et al. 2019) using the *mem* algorithm of BWA (Li and Durbin 2009) with default settings. We filtered out poorly mapped reads using the following criteria: mapping quality < 30, read not mapped in proper pairs, > 50% of the read is soft-clipped. We called genotypes as described above but with the default strand-filter of *VarScan2* (Koboldt et al. 2012). Genotypes were filtered using *VCFtools* (Danecek et al. 2011). As in the above, filters were set to allow maximum 20% missing data, maximum two alleles and not more than 20% heterozygosity per site (see above) and to remove indels and low quality genotype calls marked “filtered” by *VarScan2*. Additionally, we removed sites with a mean depth greater than 45 (twice the mean depth of all sites) and smaller than 7.5 (mean depth of all sites divided by 3). From the resulting genotype dataset, which comprised 78.976.767 nucleotide positions, we extracted a total of 2.954.526 SNPs using *VCFtools*. To verify sample identity, we performed a PCA using the *adegenet* package (Jombart et al. 2016). Based on this PCA and the RAD-seq data above, we re-assigned one *A. sagittata* mis-labeled sample (174) from one allopatric population to the other.

### Estimation of the population recombination rate

In order to take within-locus recombination into account in our ABC estimations of the demographic history, we estimated the population recombination rate R; we used the sympatric *A. sagittata* population because we had sequenced the most individuals in this population. We first phased the reads using the read-aware phasing algorithm implemented in SHAPEIT (Delaneau et al. 2013). We then used FastEPRR (Gao et al. 2016) to estimate the local recombination rate in non-overlapping 75 kbp windows along the genome, with otherwise default settings. We used the median of the distribution (R=11.5 per 75kbp window) as the per locus population recombination rate Rho=4 Ne r (i.e. Rho=0.15 per kb).

### Detection of interspecific gene flow

To investigate signatures of gene flow among non-admixed individuals, we analyzed the whole-genome resequencing (WGS) data using two four-taxon statistics: we calculated D (Durand et al. 2011) over the whole genome to detect interspecific gene flow, and we calculated f_D_ (Martin et al. 2015) in 50kbp sliding windows to analyze the heterogeneity of gene flow along the chromosomes. These tests assumed the general phylogeny: (P1,P2)(P3) outgroup; where P1 was the control population (no gene flow), P2 was the gene-flow target population and P3 was the donor population of gene flow. We used *A. androsacea* as the outgroup to determine the ancestral allelic state. We calculated f_D_ in 50 kbp sliding windows with 25 kbp overlap, following Martin et al. (2015). We skipped sites with derived allele frequency of 0 in P3, as these could bias values of f_D_. We calculated these statistics for the phylogenies shown in Figure 2.

To complement the four-taxon analysis, we applied a method based on analyses of genetic distance to locate and assign boundaries to putative introgressed fragments in individual accessions (excluding admixed individuals as in the above). This method allowed identifying introgressed fragments regardless of their frequency and their size and may be better suited to the detection of introgressions of various ages. Our reasoning was as follows: if there is no introgression, the mean genetic distance of a given accession to members of the same species should be smaller than the distance to members of other species; for introgression, the opposite is true. Thus, in 10 kbp windows along the genome, we calculated the mean intra- and interspecific genetic distance for each accession, excluding individuals from the same population for the intraspecific case, as they are likely to carry the same introgression. We added a constant of 0.0001 to all interspecific distances to avoid zero-division and removed windows with zero interspecific distance and non-extreme intraspecific distance (< 90^th^ percentile), as these could lead to spurious signals of introgression. For each window, we calculated the ratio of intraspecific and interspecific genetic distance (hereafter, distance ratio), which we used to identify introgressions.

For each accession, we identified introgressions by first removing all windows with a distance ratio smaller than or equal to 1. We then classified windows as introgressions if their distance ratio was > 2 or if both of their neighboring windows fulfilled this condition. This threshold was chosen because it seemed conservative enough to identify putative introgressed tracks. Adjacent introgression windows were connected to a single introgression. We applied several filters to the resulting set of candidate introgressions: We excluded candidate introgressions from further analysis if they contained too many repetitive elements (i.e those with mean number of genotyped positions less than the 25% quantile of the number of genotyped position in all introgressions). Furthermore, we overlapped candidate introgressions with D and f_D_ values (in 50 kbp windows) for the respective target population; for introgressions larger than 50 kbp, we calculated the mean of all windows within the introgression. We kept introgressions with values of D > 0.1 and f_D_ > 0.01. Subsequently, we removed introgressions with > 30% mean heterozygosity, as these often gave inaccurate signals: the expectation for this distance ratio is not the same as for the homozygous case (detecting heterozygous introgressions with this method would require haplotype data). To define the boundaries of each introgression accurately within each of the remaining candidate introgressions, we recalculated the genetic distances in sliding 2 kbp windows in 200 bp steps. We applied the same thresholds as described above to define our final set of introgression regions in all these windows.

Finally, we aimed to determine the likely origin of each introgression by calculating genetic distances of the introgression region between all accessions of the donor species and the target accession. We then determined the accession(s) with the minimum genetic distance to a given introgression. We standardized genetic distances by dividing them with the approximate number of genotyped sites for each pair of accessions (we used an approximation to reduce computational requirements). The approximate number was estimated by taking the mean of all 2 kbp windows (200 bp steps) within an introgression and scaling this value by the length of the introgression. If all accessions originated from the same population (i.e. sympatric or allopatric for the donor species), we inferred that the introgression likely originated from this population (or a closely related one). If donor accessions originated from multiple populations, the inferred origin was ambiguous.

Statistical analysis was performed in R (R Development Core Team 2008). Visualization was done using the ggplot2 library (Wickham 2009). To account for the introgression frequency in statistical analyses of origin and introgression size, we counted each unique introgression (identical start- and end-points) only once, regardless of its frequency. We tested whether introgression size differed among populations using a Kruskal-Wallis test followed by a pairwise Dunn-Test, implemented in the *FSA* package (Ogle et al. 2020).

### Modelling the history of interspecific gene flow

We used coalescent simulations and an ABC algorithm with random forest-based Bayesian parameter inference to determine the most likely demographic history of the two species, focusing especially on the history of interspecific gene flow (Raynal et al. 2019). Modelling was performed in two steps: i) we compared data obtained from each postulated demographic model to our observed genetic data to identify the best-fitting one and ii) we estimated the posterior distribution for each demographic parameter of the best model.

The summary statistics for the observed data were computed from the 35 fully sequenced genomes. We randomly selected 500 genomic windows of 75 kbp each, all of which were at least 150 kbp apart from one another and contained at least 20,000 genotyped sites, which allowed excluding regions that were excessively repetitive. We computed the following summary statistics using a custom python script: overall θ_w_,θ_w_ and θ_π_ within each population, Tajima’s D for each population, d_XY_ and F_ST_ between all pairs of populations, the fraction of shared, private, invariant and fixed SNPs for all pairs of populations, Patterson’s D and f_D_ (ABBA-BABA) for 8 possible phylogenies. All summary statistics, except Patterson’s D, were calculated per locus and the resulting distributions were further summarized as the mean and variance as well as the 5% and 95% quantile, resulting in a total of 236 summary statistics. To estimate the variance in model selection and parameter inference due to window choice, we assembled 100 random sets of these 500 genomic windows and considered each as one independent observation.

Simulated data for each of the five model described in Figure 3A were obtained with the software *escrm* (Sellinger et al. 2020), which is a modification of the widely used coalescent simulation software scrm (Staab et al. 2015) that accommodates selfing (we set a 90% selfing rate for all populations). Fixed parameter values and prior distributions for all demographic parameters are found in Table 1. For each model, we simulated 50,000 datasets, each consisting of five hundred 75 kbp-long loci recombining at a rate estimated from the data, as described above. Summary statistics for the observed data are computed using the same custom python script.

**Table 1:** Overview of demographic parameters (see Figure 3A) and their prior distributions or fixed values. M_Sym_ and M_Anc_ had the same prior distributions for both directions of migration. 500 loci of length 75bp were simulated assuming a per site mutation rate of 7.10^-9^ (Ossowski et al. 2010) and a population recombination rate of 0.15 per kb. Preliminary work had shown that ancestral population sizes tended to be larger, so we used a uniform distribution to improve simulation efficiency.

| Parameter | Minimum | Maximum | Distribution |
| --- | --- | --- | --- |
| $N_{N1}, N_{N2}, N_{S1}, N_{S4}$ | 500 | 100,000 | log-uniform |
| $N_{Nanc}, N_{Sanc}$ | 500 | 100,000 | Uniform |
| $N_{Anc}$ | 50,000 | 200,000 | Uniform |
| $T_N, T_S$ | 0 | 20,000 | Uniform |
| $T_{Anc}$ | 500,000 | 1,500,000 | Uniform |
| $M_{Sym}, M_{Anc}$ | $1e-11$ | $1e-3$ | log-uniform |
| $M_N, M_S$ | $1e-11$ | $1e-1$ | log-uniform |

### Model choice procedure

In the first step, we specified five different demographic models, all of which differed in their mode of intra- and interspecific gene flow (Figure 3A). In the first model, we assumed no migration at all, either intra- or interspecific. In the second model, interspecific migration stopped 100,000 generations after the species split. In the third model, ancestral interspecific migration continued until the first intraspecific population-split. The fourth model was an extension of the third model that additionally allowed migration between the sympatric populations, after populations in both species had split.

In the fifth model, interspecific migration continued throughout the history of the species, allowing a change in intensity after the intraspecific population had split in both species. All interspecific migration rates were allowed to be asymmetrical, while symmetric intraspecific migration was assumed in all models (except model # 1, which assumed no migration at all). Population sizes were constant for each population but were allowed to change at all population split points (Table 1).

We used *abcrf,* an R package designed for ABC-based demographic modelling based on random forests, to choose the model that best fit our data and to infer their parameters (Csilléry et al. 2015, Marin et al. 2019; Raynal et al. 2019). We trained a random forest classifier to distinguish between the five models based on a set of summary statistics, i.e., the model was the dependent variable and the 236 summary statistics were independent variables. We trained the random forest algorithm on 50,000 simulated datasets for each of the five models drawing from the priors in Table 1 and using *abcrf* default parameters, except that parameter “lda” was set to false, disabling linear discriminant analysis. To estimate the classification error for model choice, we used the so-called out-of-bag method implemented in *abcrf*. Due to bootstrap aggregating, each sample was used only to build a subset of trees in the random forest, and the remaining trees could be used to predict the best model for this sample. The results were represented as a confusion matrix showing how frequently the predicted model matches (or does not match) the true model (Figure 3 B). All samples were expected to fall on the diagonal of such a matrix. We then used the trained classifier to assign the most likely demographic model to each of the 100 observed samples (see above). The model that was most frequently selected as the best model for the observed samples was selected as the model fitting our data best.

We further quantified the goodness-of-fit for the best model using a method implemented in the R package *abc* (Katalin et al. 2015), which is based on a rejection algorithm. Briefly, the normalized distance of summary statistics between the observed and all 50,000 simulated datasets was calculated; then the median of the lowest 1% of distances was extracted (the rejection step). The same procedure was performed for 1,000 simulated datasets under this model, drawing parameter values from the prior distributions of distances (so-called pseudo-observed distances). If the model fit is good, the observed distance (1% best-fitting simulations to the observed data) should lie well within the distribution of pseudo-observed distances (1% best simulations fit each of the pseudo-observed dataset). We conducted this analysis for all 100 observed datasets. For each dataset, a p-value was calculated as the fraction of pseudo-observed distances, which are higher than the observed distance.

### Estimation of demographic parameters

After identifying the best-fitting demographic model, we estimated the posterior distributions of the different parameters. Demographic parameters with log-uniform prior distributions were log-transformed prior to training to make them uniform. To reduce computational load, we first selected a subset of the most informative summary statistics for each parameter; we then trained a random forest for each demographic parameter based on 115,000 simulated datasets and extracted the top ten most important summary statistics. This approach reduced the 236 summary statistics to a set of 111 unique summary statistics for training the full models.

The accuracy of random-forest predictions can be greatly increased by tuning training parameters. We tuned the following training parameters by training random forests using all possible combinations: the number of variables to possibly split in each node (mtry; search space: 11, 22, 44, 66, 88); and the minimal node size (search space: 5, 10, 30, 50). This training was conducted for a subset of representative demographic parameters (N_N1_, N_Nanc_, T_N_, M_SN,anc_, M_S,_ M_SN,sym_). We plotted the out-of-bag prediction error for each combination of training and demographic parameters (Figure S11). Based on these results, we took mtry=44 as the optimal number of randomly sampled variables, as higher values strongly increased training time while increasing accuracy only minimally. The influence of minimal node size was generally small, so we kept it at the default value of 5.

Finally, we trained a random forest for each parameter on 359,000 simulated datasets, keeping 1,000 pseudo-observed datasets out of the training dataset to use as a test. We quantified the estimation error for each model parameter by computing the parameter estimates for each pseudo-observed dataset and calculating R² between true and estimated values. Additionally, we computed the root-mean-square error (RMSE) per model parameter as: 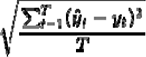, where T is the number of pseudo-observed datasets, y_t_ is the true value and ŷ_t_ is the estimated value. To make RMSE more comparable among parameters, we further converted RMSE to percentages (%RMSE) by scaling it to the prior range of each parameter (Figure S8).

To calculate parameter estimates for the observed data, we first randomly selected one of the 100 genomic sub-samples (observed samples) that was previously classified as fitting the best model For this dataset, point estimate, standard deviation, posterior mode and the 5% and 95% quantiles of the posterior distributions were determined for each demographic parameter. Additionally, we determined point estimates for the remaining 99 observed datasets to assess the variation of demographic parameters depending on the chosen genomic sub-sample.

To complement *abcrf,* which estimates a posterior distribution, we also used *XGBoost* (Chen et al. 2020), a widely used tree-based machine-learning method that gives only a point estimate but often achieves greater accuracy than random forests. We included all summary statistics in XGBoost models. We split the 50 000 simulated datasets into a test dataset of 20,000 samples and a training dataset with the remaining 30,000 samples. Tree depth (dp; values: 3, 6, 9) and minimum child weight (values: 5, 10), are the two training parameters that control the complexity of the trees. We tuned these parameters for the same demographic parameters as described above, chooseing those that minimized RMSE (Figure S12). The algorithm was stopped if the test statistic did not improve for 100 rounds. For tree depth, we chose six as the optimal value for training models for T_S_, T_N_, N_Nanc_ and N_Sanc_, and nine for all other model parameters. We chose 10 as the optimal value for minimum child weight. Using these settings, we trained an XGBoost model for each demographic parameter, again stopping after 100 rounds if there was no improvement. We assessed accuracy using the test dataset with 20,000 simulated observations, as described for *abcrf*. We then performed the estimation of the demographic parameters for the 100 genomic sub-samples forming the observed datasets. Posterior distributions are shown in Figure S13.

## ACKNOWLEDGMENTs

This research was funded by the European Research Council (ERC) through the “AdaptoSCOPE” grant 648617, and by the Deutsche Forschungsgemeinschaft through DFG Grant ME2742/13-1. HD acknowledges funding by the International Max Planck Research School. We thank Dr. Markus Stetter and Dr. Gregor Schmitz and Prof. N. Hölzel for helpful discussions. Sequence data are available at ENA under ERS5040446-ERS5040483. Markdown scripts are available as supplementary information.

## SUPPLEMENTARY MATERIAL

Suppl_Fig.pdf: Supplementary figures S1-S13 with legends.

RADseq.html: R Markdown of RAD-seq genotyping script and analyses.

Introg_Det_final.html: R Markdown of Introgression detection in whole genome data Demography_estimation_final2.html: R markdown for ABC modeling Suppl_Tables: Table S1 Description of genotypes included in the RAD-seq analysis, Table S2: Description of genotypes chosen for whole-genome sequencing –Table S3: list of introgressed fragments identified in the genomes of individuals described in Table S2. Table S4: Summary statistics describing the 100 observed datasets. Table S5: ABCRF parameter distribution and XGBoost point estimate for the representative observed dataset shown in Fig. 4. Table S6: Variation in parameter point estimates for the 100 observed datasets.

