## Supplementary material for "Approximate Bayesian computation untangles signatures of contemporary and historical hybridization between two endangered species": Suppl figures

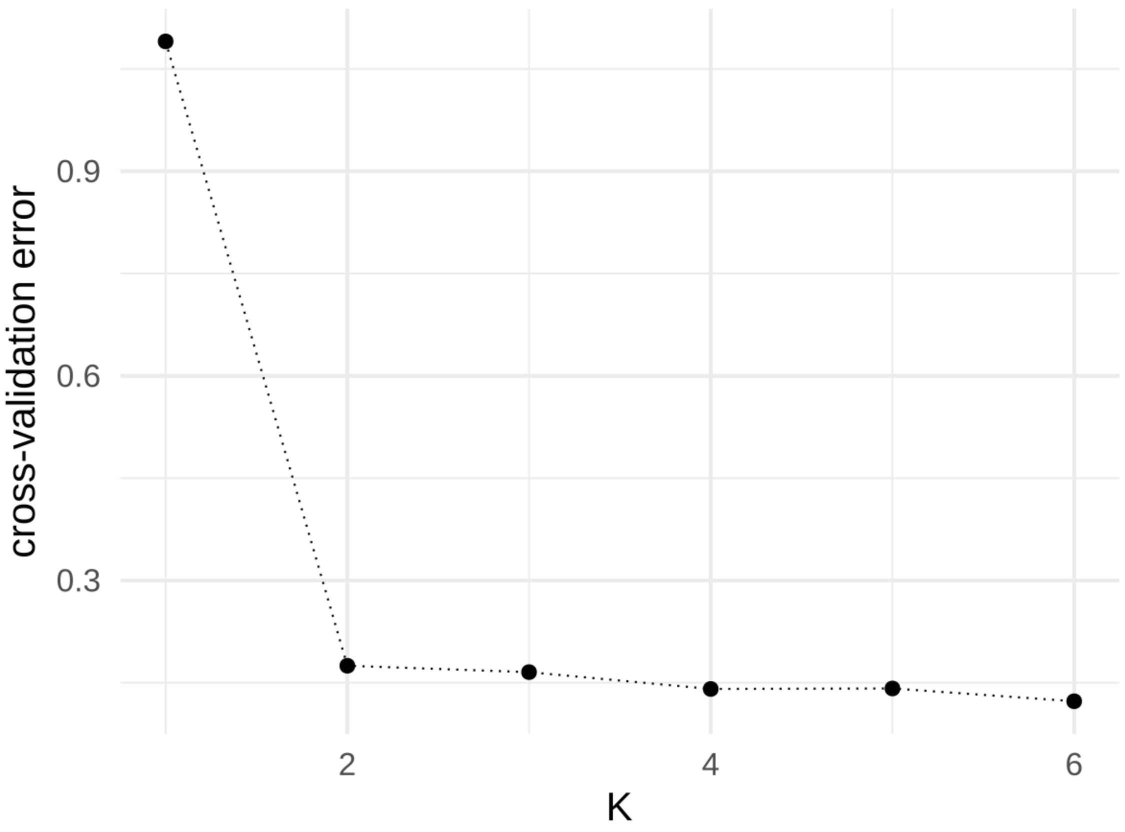


*Figure S 1: Cross-validation error of the Admixture analysis with K ranging from 1 to 6.*

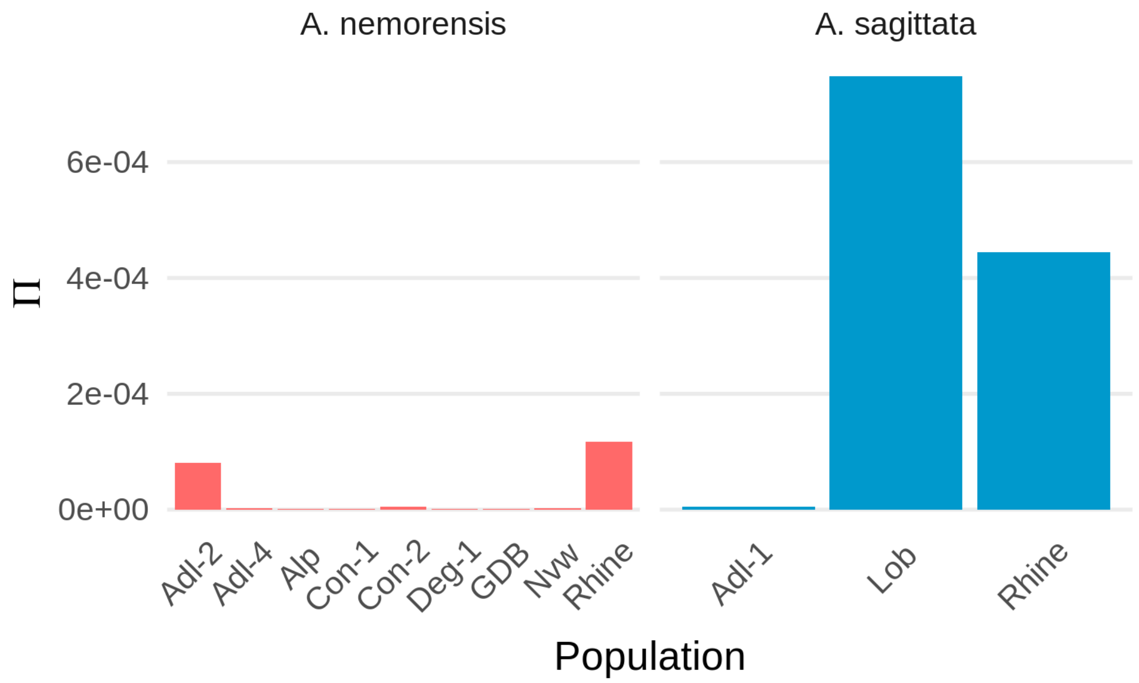


*Figure S 2: Genetic diversity within each population of the two species. Genetic diversity was measured as the average number of pairwise differences per base pair.*

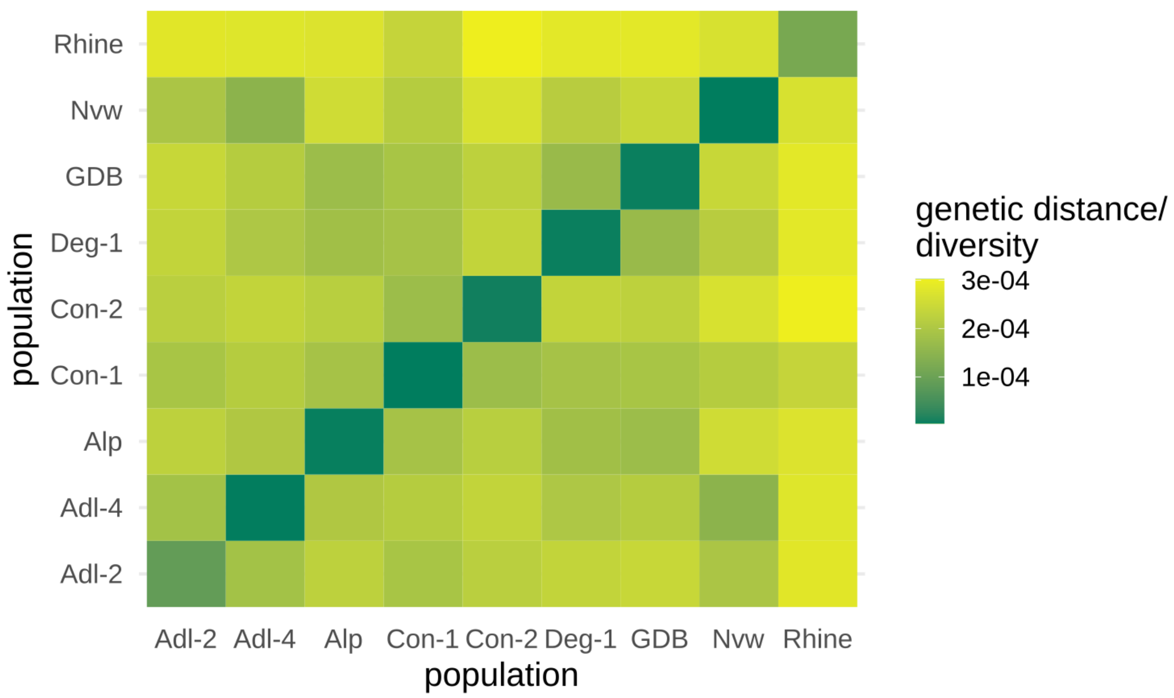


*Figure S 3: Pairwise genetic distance among populations of A. nemorensis. Genetic distance was measured as the average number of pairwise differences per base pair. The diagonal line represents the diversity within each population.*

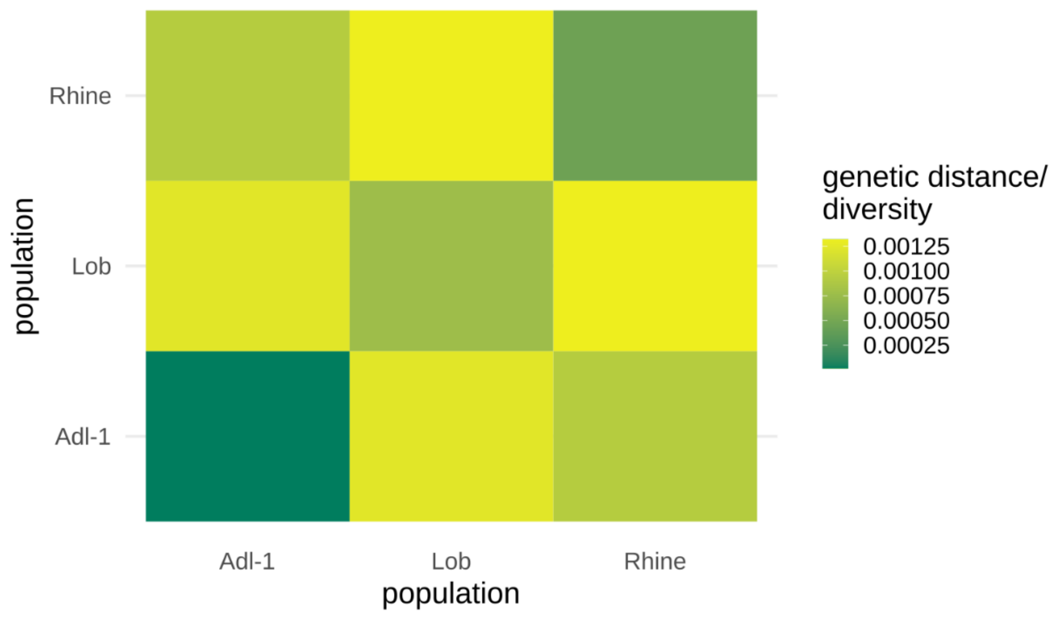


*Figure S 4: Pairwise genetic distance among populations of A. sagittata. Genetic distance was measured as the average number of pairwise differences per base pair. The diagonal line represents the diversity within each population.*

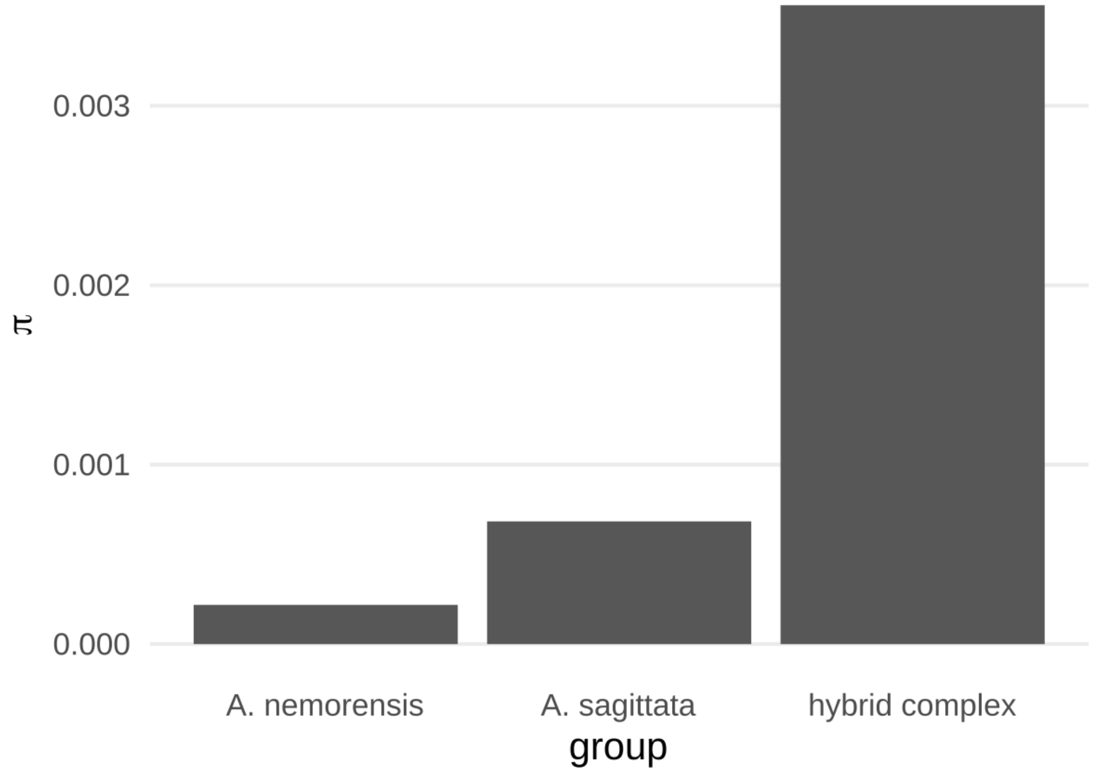


*Figure S 5: Genetic diversity within each species and within the hybrid complex, which contains the whole sympatric population (parental species and hybrids). Genetic diversity was measured as the average number of pairwise differences per base pair.*

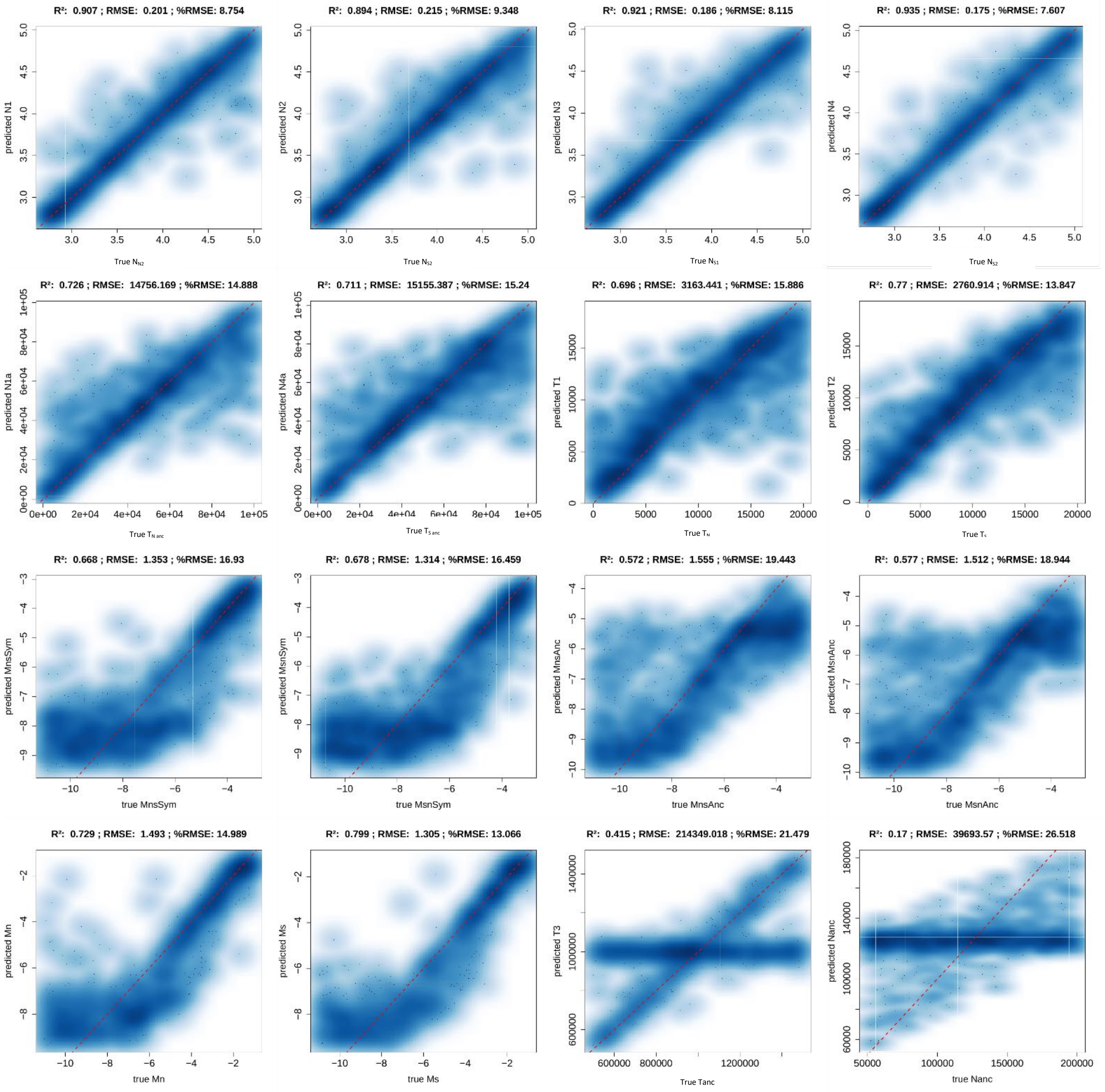


*Figure S 6: Correlation plots of true and predicted parameter values for 1000 simulated samples per parameter. The x-axis shows the true value and the y-axis the value predicted by the abcrf model. The dashed red line represents the theorethical 1:1 relationship. Parameter names deviate from the main text: N1 – Effective population size of allopatric A. nemorensis populations $N_{N1}$, N2 – Effective population of sympatric A. nemorensis population $N_{N2}$, N3 – Effective population of sympatric A. sagittata population $N_{S1}$, N4 – Effective population size of allopatric A. sagittata population $N_{S2}$, N1a – Ancestral population size of A. nemorensis $N_{Nanc}$, N4a –Ancestral population size of A. sagittata $N_{Sanc}$, T1 – time elapsed since split of A. nemorensis population $T_N$, T2 – Time elapsed since split of A. sagittata populations $T_S$, T3 – time of speciation $T_{Anc}$, N3 – ancestral population size prior to speciation $N_{Anc}$*

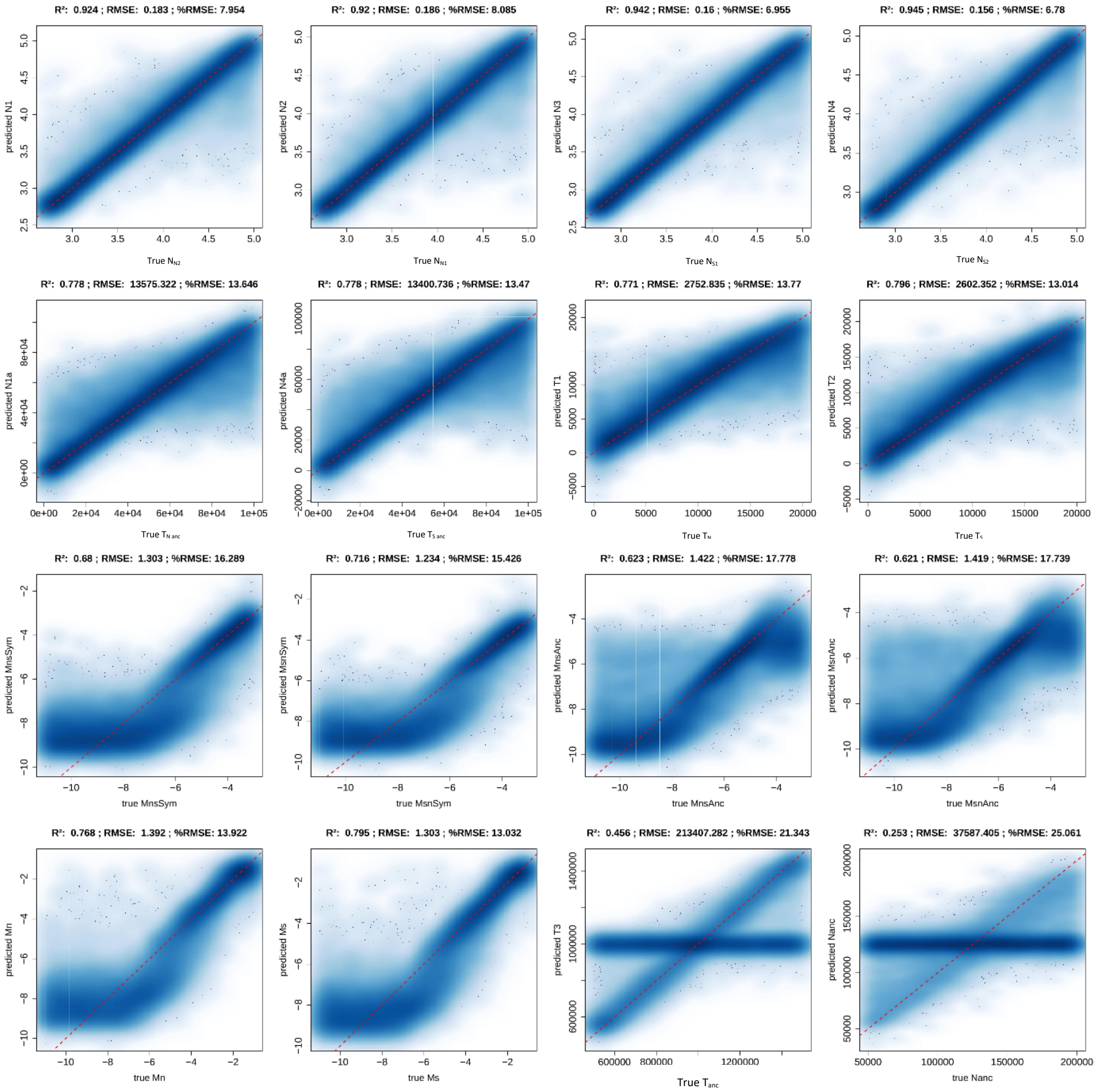


*Figure S 7: Correlation plots of true and predicted parameter values for 20,000 simulated samples per parameter. The x-axis shows the true value and the y-axis the value predicted by the XGBoost model. The dashed red line represents the theorethical 1:1 relationship. Parameter names deviate from the main text: N1 – Effective population size of allopatric A. nemorensis populations $N_{N1}$, N2 – Effective population of sympatric A. nemorensis population $N_{N2}$, N3 – Effective population of sympatric A. sagittata population $N_{S1}$, N4 – Effective population size of allopatric A. sagittata population $N_{S2}$, N1a – Ancestral population size of A. nemorensis $N_{Nanc}$, N4a –Ancestral population size of A. sagittata $N_{Sanc}$, T1 – time elapsed since split of A. nemorensis population $T_N$, T2 – Time elapsed since split of A. sagittata populations $T_S$, T3 –time of speciation $T_{Anc}$, N3 – ancestral population size prior to speciation $N_{Anc}$*

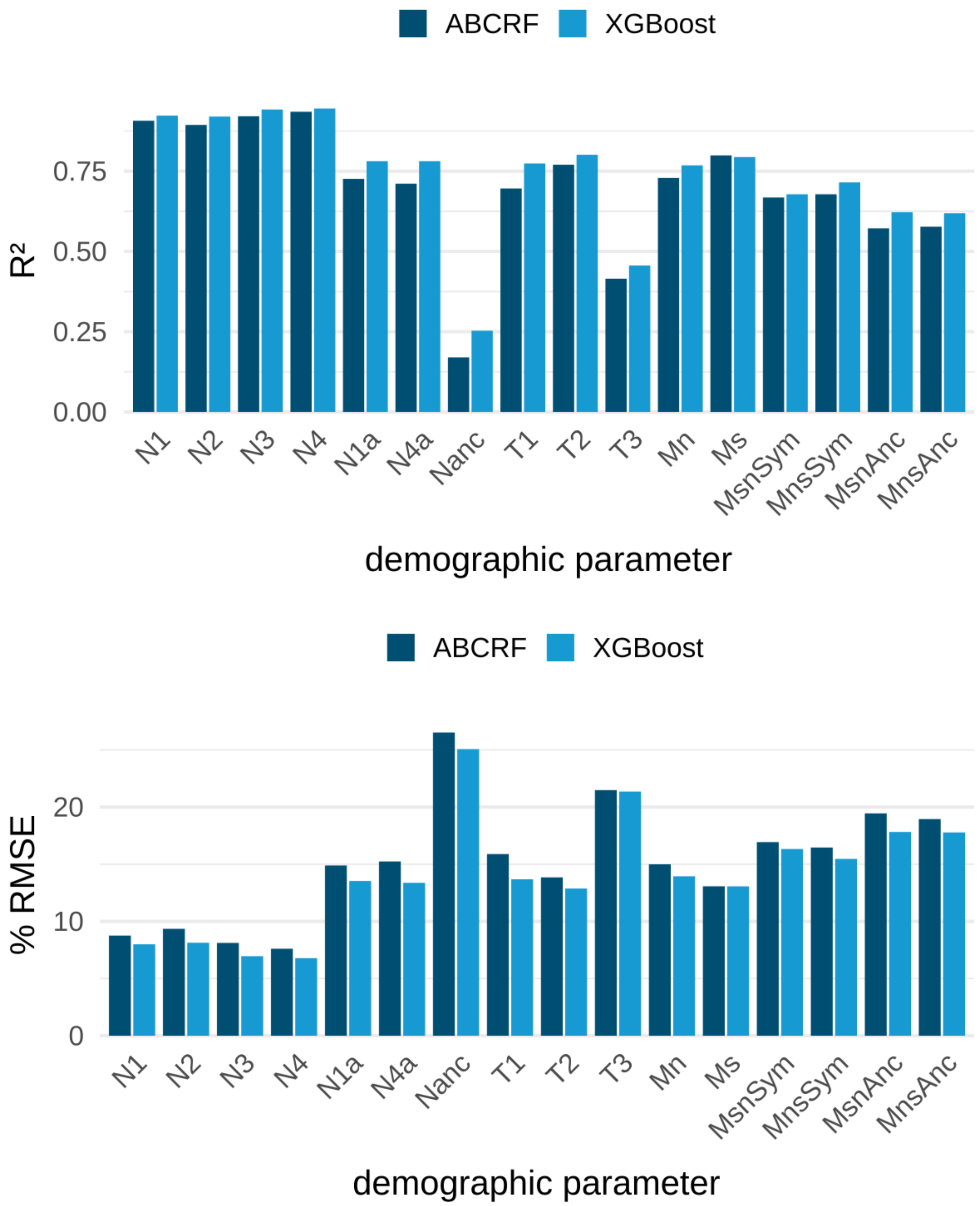


*Figure S 8: Comparison of accuracy statistics of ABCRF and XGBoost models. $R^2$ and RMSE were calculated between true and predicted values of each demographic parameter (see also Figure S1). RMSE was scaled by the total prior range of each parameter. A higher $R^2$ and a lower RMSE indicate higher prediction accuracy.*

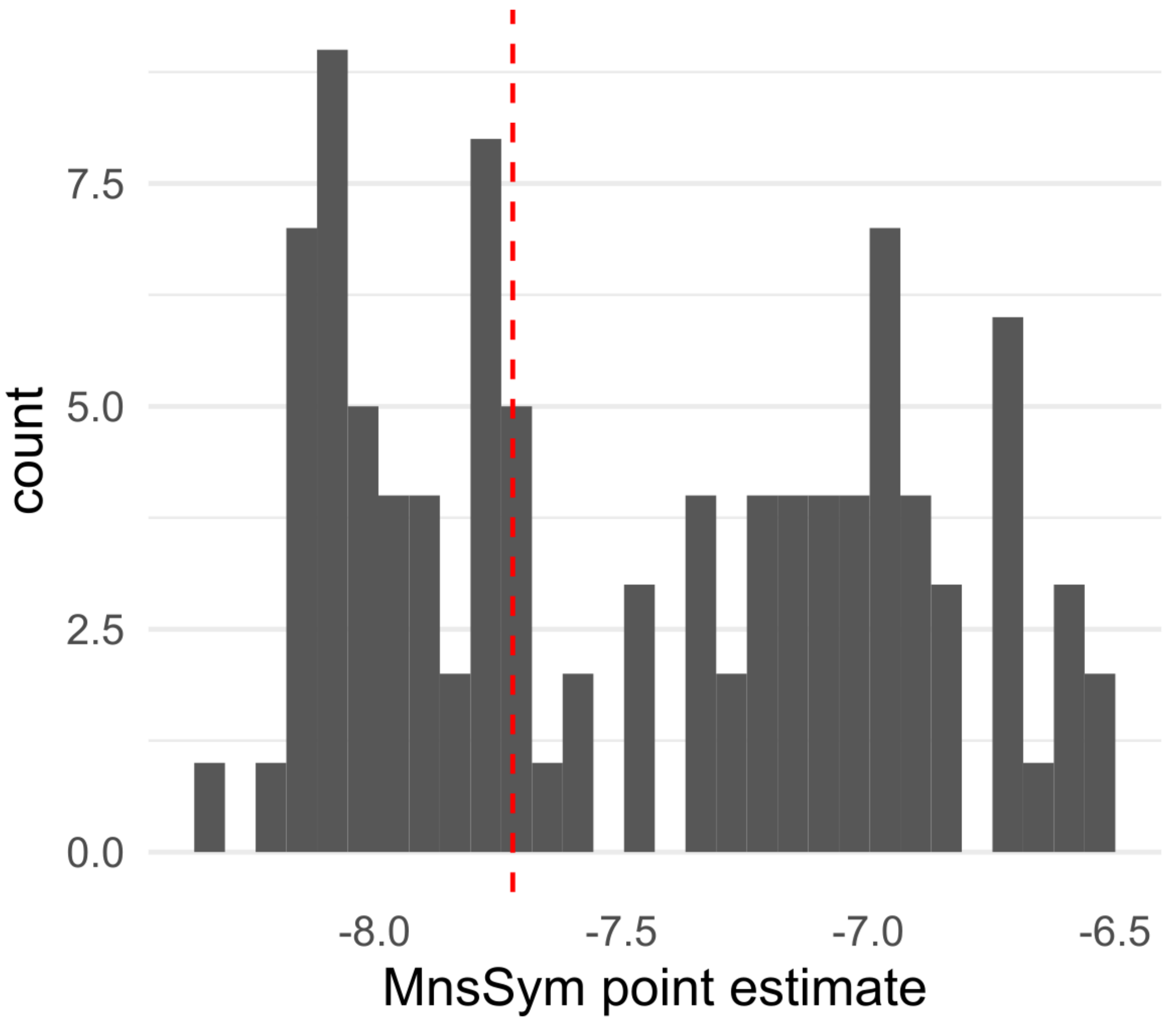


*Figure S 9: Distribution of point estimates for the sympatric migration rate from A. nemorensis to A. sagittata estimated for 100 observed samples. The dashed red line represents the sample described in the main text. Migration rates were log10-transformed.*

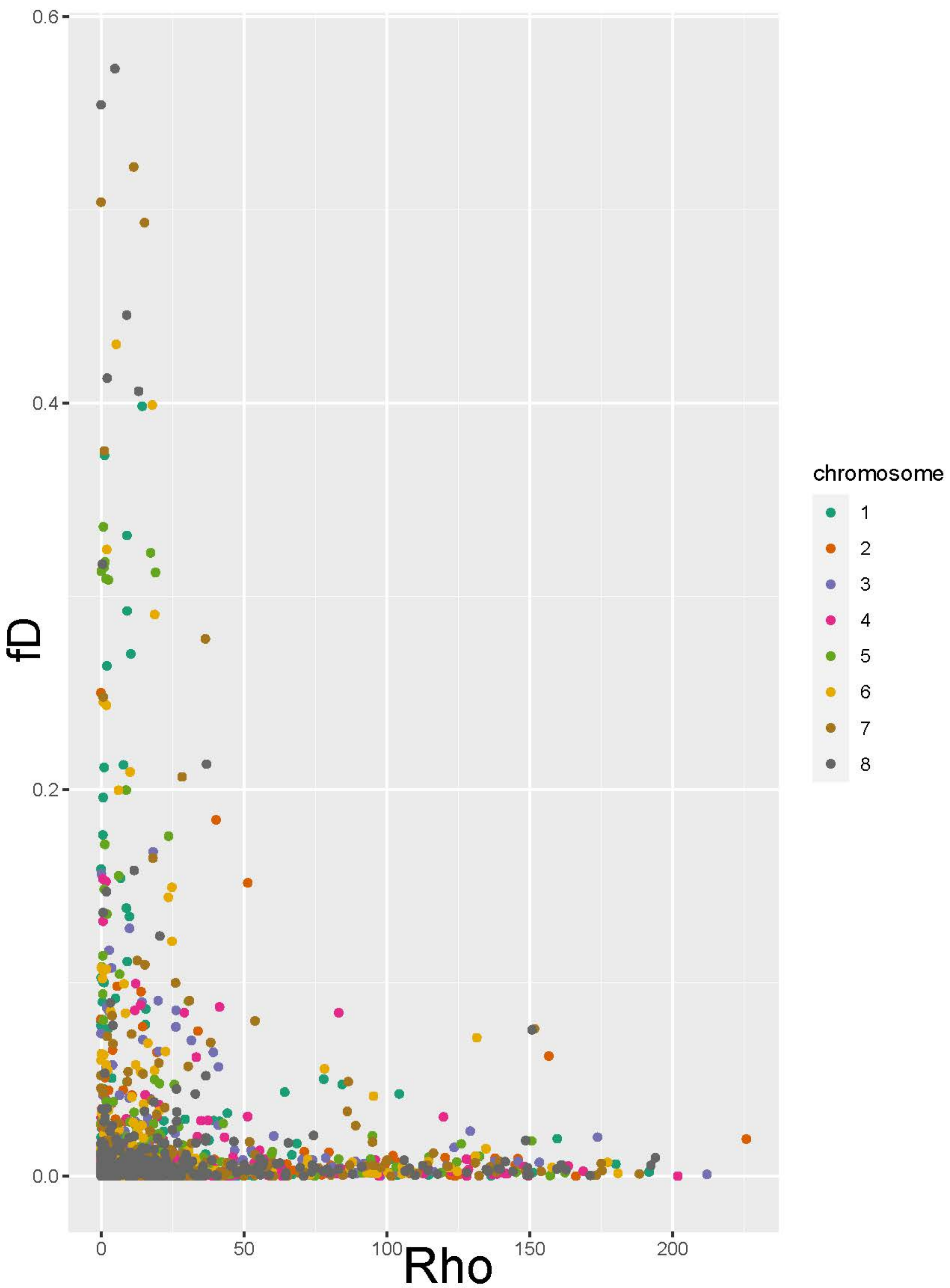


Figure S10: Mean fD of each 75kb fragment for which Rho was estimated. The relationship is weakly negative but significant (Rho= -0.06, p<0.005), showing that the strongest signals of introgression are found in regions of low recombination.

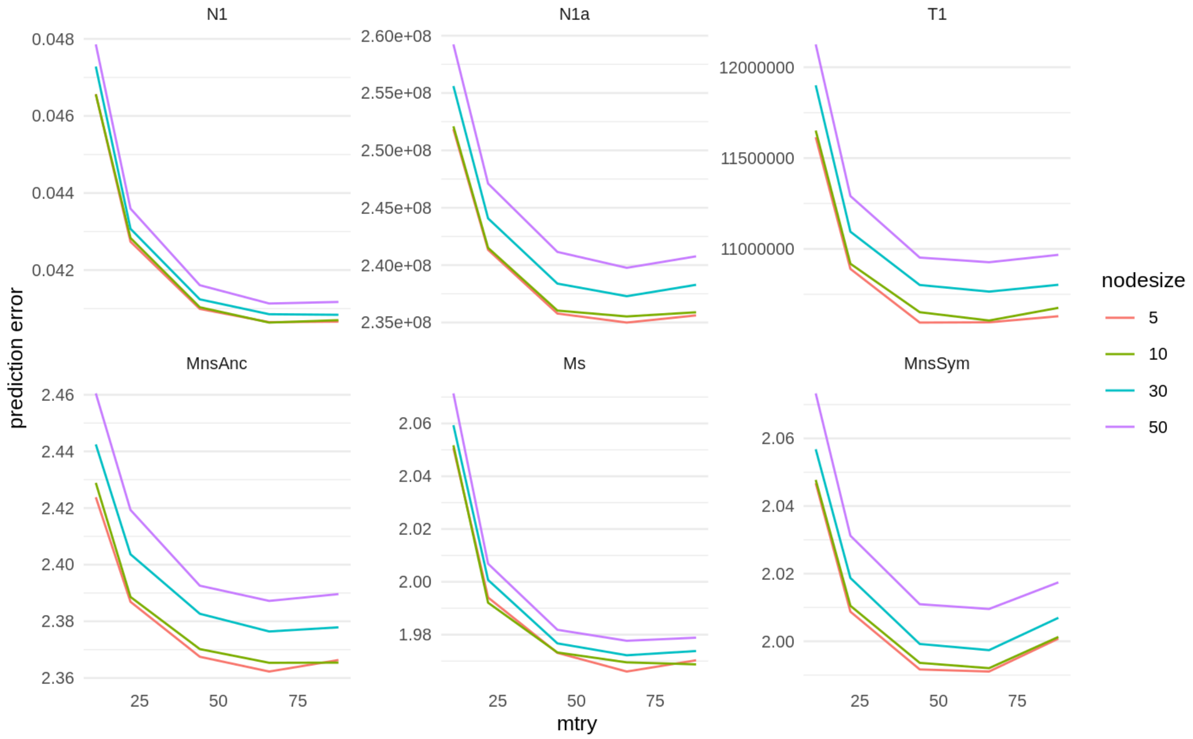


*Figure S 11: Prediction error of abcrf models trained with different combinations of the mtry and nodesize parameters. Predction error is the mean square error. Some parameter names deviate from the main text: N1 – $N_{N1}$, N1a – $N_{Nanc}$, T1 – $T_N$, T2 – $T_S$*

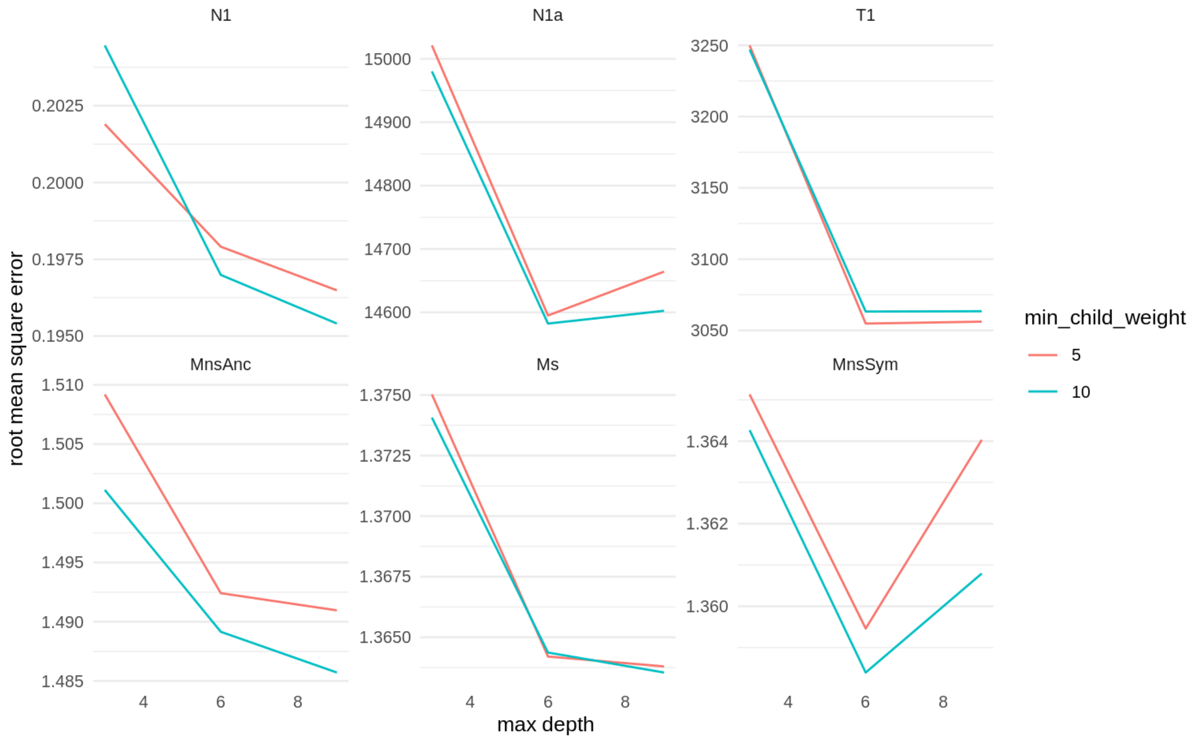


*Figure S 12: Prediction error of XGBoost models trained with different combinations of the max depth and min_child_weight parameters. Some parameter names deviate from the main text: N1 – $N_{N1}$, N1a – $N_{Nanc}$, T1 – $T_N$, T2 – $T_S$*

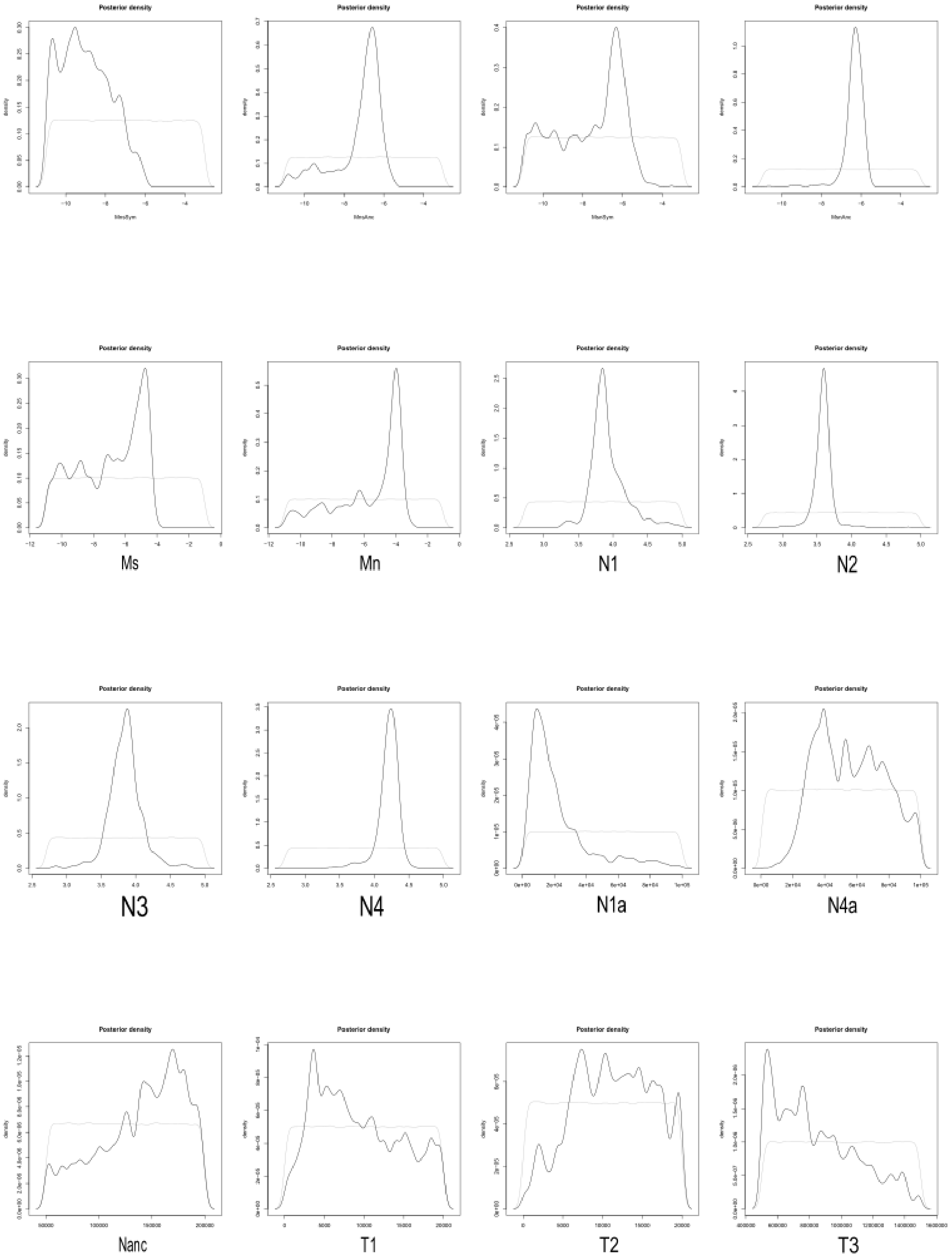


*Figure S 13: Posterior distributions of parameters estimated with ABCrf. Some parameter names deviate from the main text: N1 – $N_{N1}$, N1a – $N_{Nanc}$, T1 – $T_N$, T2 – $T_S$. Light grey line shows the prior, dark grey line the posterior distribution.*
